# Mapping the Temporal Landscape of Breast Cancer Using Epigenetic Entropy

**DOI:** 10.1101/2024.10.11.617056

**Authors:** Daniel L. Monyak, Shannon T. Holloway, Graham J. Gumbert, Lars J. Grimm, E. Shelley Hwang, Jeffrey R. Marks, Darryl Shibata, Marc D. Ryser

## Abstract

Although generally unknown, the age of a newly diagnosed tumor encodes valuable etiologic and prognostic information. Here, we estimate the age of breast cancers, defined as the time from the start of growth to detection, using a measure of epigenetic entropy derived from genome-wide methylation arrays. Based on an ensemble of neutrally fluctuating CpG (fCpG) sites, this stochastic epigenetic clock differs from conventional clocks that measure age-related increases in methylation. We show that younger tumors exhibit hallmarks of aggressiveness, such as increased proliferation and genomic instability, whereas older tumors are characterized by elevated immune infiltration, indicative of enhanced immune surveillance. These findings suggest that the clock captures a tumor’s effective growth rate resulting from the evolutionary-ecological competition between intrinsic growth potential and external systemic pressures. Because of the clock’s ability to delineate old and stable from young and aggressive tumors, it has potential applications in risk stratification of early-stage breast cancers and guiding early detection efforts.

## INTRODUCTION

When a woman is diagnosed with breast cancer, it is generally not possible to ascertain how long the tumor has been growing. Yet knowledge about a tumor’s age at diagnosis could provide important prognostic clues: older indolent tumors that are less likely to progress may require less invasive treatment, whereas younger fast-growing tumors require more urgent and aggressive treatment. While there is a rich literature on the estimation of the mean sojourn time of breast cancer^1–5^—the average time tumors spend in a detectable but asymptomatic, pre-clinical state—there is a paucity of tools to assess the age of individual tumors at the time of detection.

Epigenetic clocks provide a promising approach to estimate individual tumor age. Originally developed to quantify the biologic aging process in humans, epigenetic clocks leverage specific patterns of DNA methylation that are strongly correlated with biologic tissue age.^6,7^ Broadly, these clocks focus on the methylation status of CpG sites across the genome that are unmethylated at birth and become methylated with increasing tissue age due to the accumulation of stochastic replication errors. Due to a low error rate, such clocks are well suited to resolve processes that evolve over time scales on the order of decades, such as normal tissue aging and pre-neoplastic transformations.^8–12^ They are, however, less useful for estimating the age of invasive cancers, which take mere months to years to become symptomatic or reach a screen detectable size.

In contrast to “one-way” clocks that measure age-related increases of methylation, “two-way” epigenetic clocks leverage CpG sites that fluctuate between the unmethylated and methylated states on a relatively fast time scale (**Figure 1A**). Originally introduced in the context of homeostatic intestinal stem cell dynamics,^13^ such stochastic epigenetic clocks have also been applied to hematologic malignancies.^14^ Here we follow similar design principles to develop a breast cancer-specific two-way epigenetic clock to measure individual tumor age at diagnosis based on average methylation levels (*β*-values) of select CpG sites included in standard methylation arrays.

**Figure 1.**
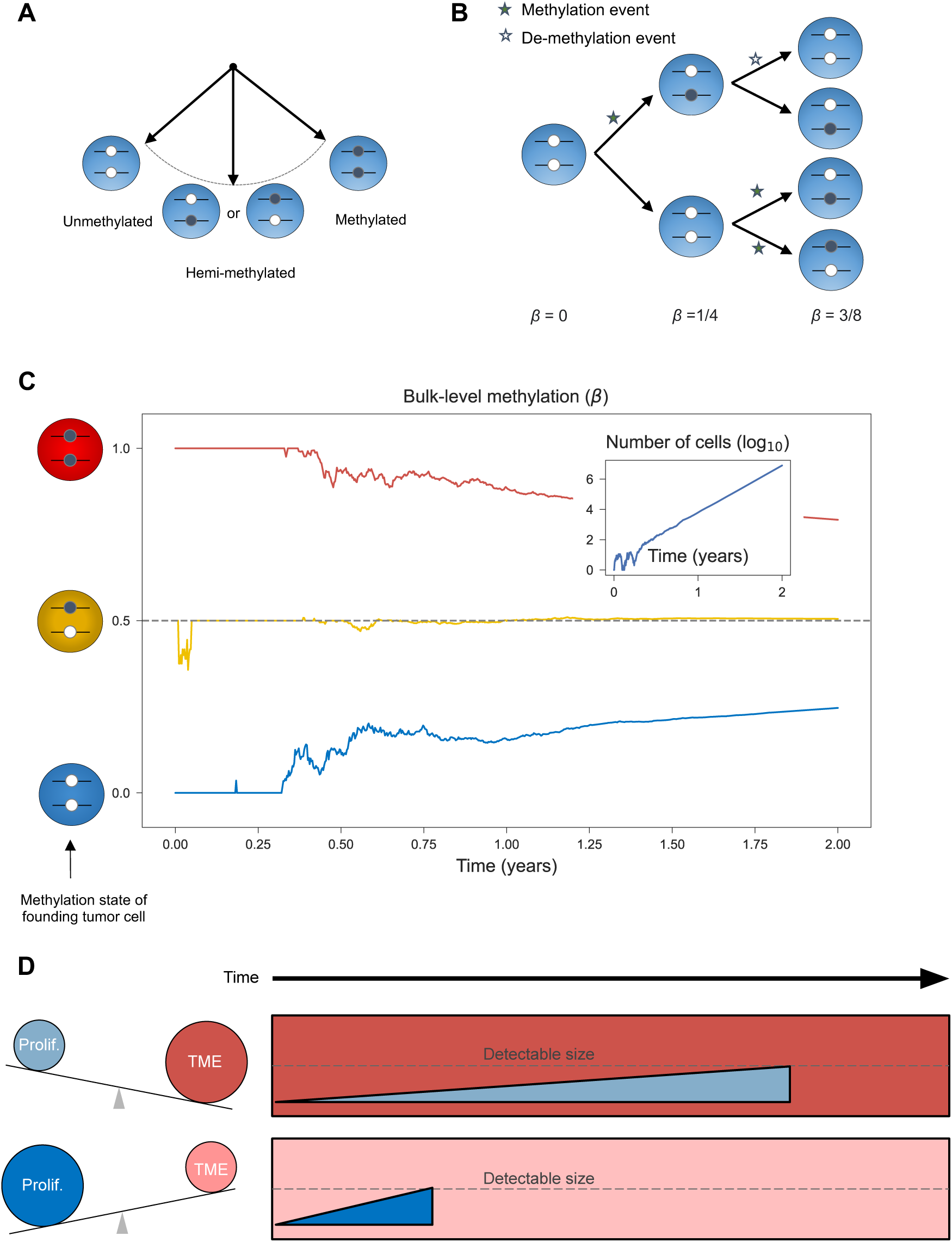
Fluctuating CpG sites (fCpG) and modulators of tumor age. **(A)** Balanced fCpGs stochastically oscillate between the unmethylated, hemi-methylated, and methylated states. **(B)** During tumor cell division, methylation and de-methylation events occur stochastically. As illustrated for the first three generations of an exponential tumor expansion, the average methylation status (*β*-value) evolves over time. **(C)** In silico modeling of the fCpG dynamics, based on a simple birth-death model of tumorigenesis. The average methylation value (*β*) is simulated for three independent fCpGs, whose initial states in the first tumor cell are fully methylated (red), hemi-methylated (yellow), and unmethylated (blue), respectively. Insert: the number of tumor cells over time. Simulation parameters: cell proliferation rate *α* = 0.17 divisions/day; cell death rate *λ* = 0.15 deaths/day; (de-)methylation rate *μ* = 0.002 flips/division. See Methods for details. **(D)** The age of a tumor at detection is modulated by relative strengths of intrinsic proliferation and growth-suppression induced by the immune tumor microenvironment (TME); tumors will reach a detectable size faster when proliferation is greater and/or the TME is weaker.

The proposed stochastic clock measures the entropy of an ensemble of fluctuating CpG (fCpG) sites. In the tumor’s most recent common ancestor cell, each fCpG was either unmethylated (*β* = 0), hemi-methylated (*β* = 0.5), or methylated (*β* = 1). As the tumor expands, replication errors produce a mixture of cells with different methylation states (**Figure 1B**), thus progressively increasing the tumor’s epigenetic entropy.^15^ In the special case of unbiased fCpG sites—whose methylation and demethylation rates are in balance—the bulk-level methylation converges to *β* = 0.5 with increasing tumor age, regardless of the first tumor cell’s state (**Figure 1C**). Thus, by measuring the distribution of unbiased fCpG sites, we can derive an estimate of the age of a given tumor cell population relative to the start of the most recent clonal expansion.

The combination of tumor-specific age estimates and gene expression profiles further provides a unique opportunity to characterize the evolutionary and ecological pressures that shape the temporal landscape of breast cancer. Notably, aggressive tumors that evolve in a weakly suppressive immune microenvironment are expected to reach a detectable size faster than indolent, slow-growing tumors in a strongly suppressive immune microenvironment (**Figure 1D**).

The manuscript is structured as follows. Combining DNA methylation and gene expression data from several hundred breast cancer and normal breast tissue samples, we first identify a set of unbiased fCpG sites and introduce the epigenetic clock index as a proxy measure of tumor mitotic age. We then evaluate the face validity of the index by examining its relationship with established prognostic markers, and we combine methylation and gene expression data to identify tumor- and microenvironment-specific factors that modulate tumor age. Finally, we validate key properties of the clock index in independent cohorts of patients with paired primary-metastasis samples, and we derive quantitative estimates of individual breast cancers’ mitotic and calendar ages.

## RESULTS

### Selection of unbiased fCpG sites

To identify a set of unbiased fCpG sites in breast cancer, we used 450K methylation array data from 634 invasive breast cancers in The Cancer Genome Atlas^16^ (TCGA) and 79 normal breast tissue samples.^17^ Using a three-step selection process, we identified an ensemble of fCpG sites with balanced (de-)methylation rates as follows.

In the first step, we eliminated regulatory and genic loci due to their increased likelihood of being under selection during tumor growth. This choice contrasts with the majority of cancer epigenomic studies focused on functional CpG sites located in or around promoter regions.^17–19^ Next, we identified CpG sites with an average *β*-value close to 0.5 in both normal breast tissue and breast cancers, thus excluding sites with an inherent bias toward methylation or de-methylation, and sites that are subject to systematic selection during homeostasis and/or tumorigenesis (**Figure 2A**). In the third step, we ordered the set of unbiased CpG sites by between-tumor variability and included only the 500 most fluctuating sites in the final clock set of unbiased fCpGs (**Figure 2B**). Importantly, this final step excludes non-informative sites that either do not fluctuate at all (i.e., imprinted hemi-methylated state) or fluctuate too fast (i.e., steady-state methylation of *β* ≈ 0.5 reached on time scales much shorter than the average mitotic tumor age at diagnosis).

**Figure 2.**
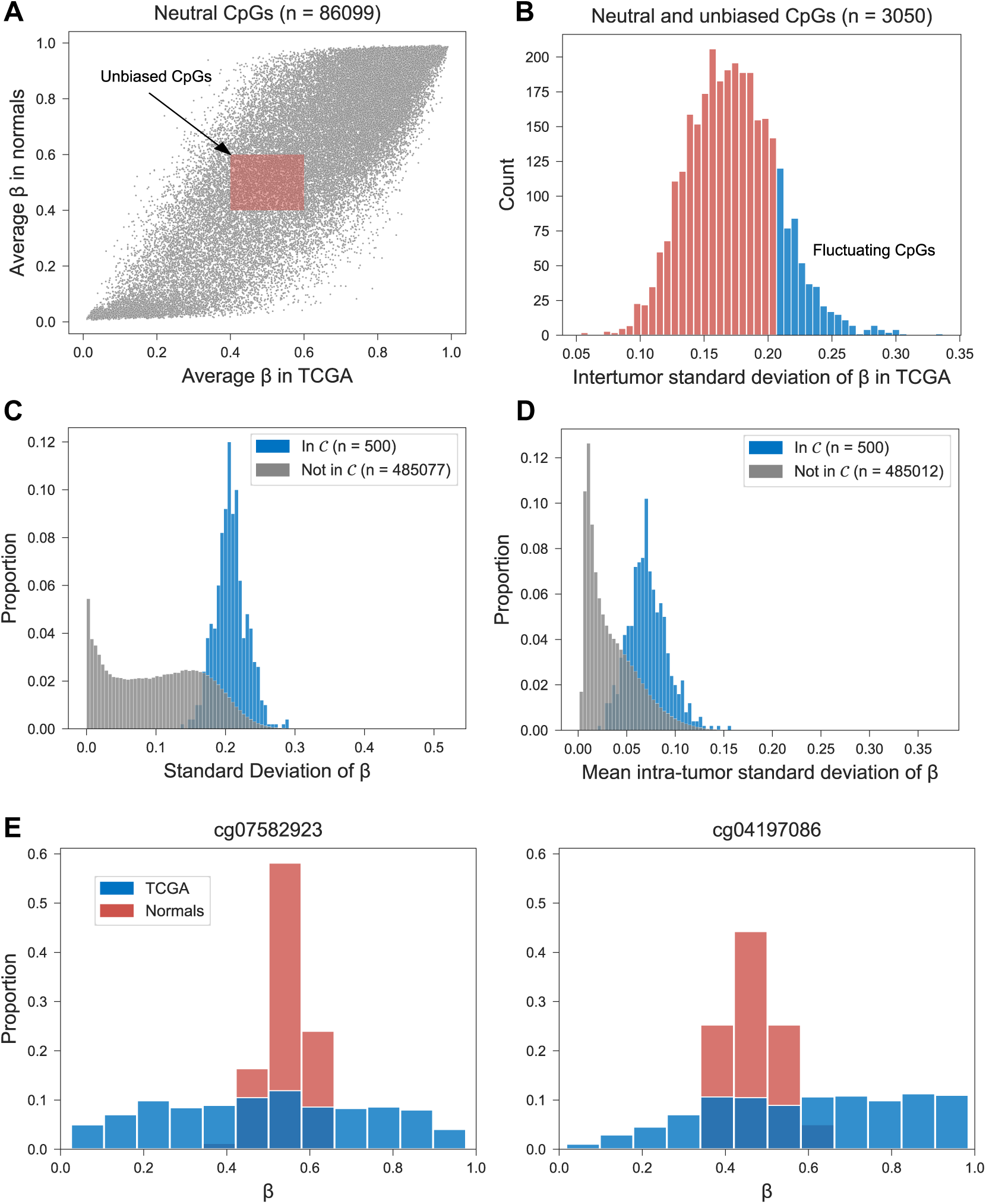
Selection and validation of unbiased fluctuating CpG (fCpG) sites. **(A)** Starting with CpG sites that are non-regulatory and non-genic (gray dots), sites with an average methylation value *β* between 0.4 and 0.6 in both breast cancers (N=634, TCGA cohort) and normal samples (N=79, Normal cohort) were considered unbiased (red shaded rectangle). **(B)** Sites were ranked by their inter-tumor standard deviation, and the top 500 were included the clock set 𝒞. **(C)** In a separate cohort of breast cancers (N=146, Lund cohort), the inter-tumor standard deviation of *β* was higher for fCpG sites in the clock set 𝒞 (median: 0.206) compared to CpGs not included in 𝒞 (median: 0.102; P=1×10^−238^, Wilcoxon rank-sum test). **(D)** In a small cohort of patients (N=5) with multiple primary samples (3-5 per patient), the intra-tumor standard deviation of *β* was higher for CpG sites in the clock set 𝒞 (median: 0.070), compared to CpGs not included in 𝒞 (median: 0.026; P=6×10^−179^, Wilcoxon rank-sum test). **(E)** Distribution of the *β*-values of two fCpG sites in the clock set 𝒞, across breast cancers (N=634, TCGA cohort) and normal breast samples (N=79, Normal cohort).

Next, we sought to validate the unbiased and fluctuating nature of the clock set in two independent cohorts. In a cohort of 146 breast cancer patients (Lund cohort),^20^ we found significantly higher inter-tumor variability in *β*-values among the CpG sites in the clock set, as compared to the CpG sites not included in the clock set (**Figure 2C**). Similarly, in a small cohort of 5 patients with multiple samples from their primary tumors,^21^ we found elevated intra-tumor variability in clock set vs non-clock set sites (**Figure 2D**). Together, these patterns corroborate the unbiased and fluctuating nature of the clock set of CpG sites.

Interestingly, the fCpG sites in the clock set were more tightly concentrated around *β* = 0.5 in normal breast tissue, as compared to breast cancers (**Figure 2E**), with a median standard deviation of *β*-values in normal samples of 0.09, compared to 0.21 in tumors (P=2×10^−45^, Wilcoxon rank sum test). Consistent with the underlying dynamic model of the clock (**Figure 1A**), this suggests that over decades of breast development and maintenance, the fCpGs had converged to the stationary methylation state of *β* = 0.5 (**Figure 1C**).

### Epigenetic clock index

At the level of individual tumors, the 500 fCpG sites in the clock set exhibited primarily unimodal or bimodal distributions of *β*-values (**Figure 3A**). We explored how these tumor-specific distributions of *β*-values could be used to estimate tumor mitotic age. In the founding tumor cell, each fCpG starts in either the unmethylated (*β* = 0), hemi-methylated (*β* = 0.5), or methylated (*β* = 1) state (**Figure 1A**). Although the trajectories of individual sites are subject to stochastic fluctuations (**Figure 1C**), an ensemble of sites starting in the same initial configuration collectively drift toward the steady state of *β* = 0.5 (**Figure 3B**).

**Figure 3.**
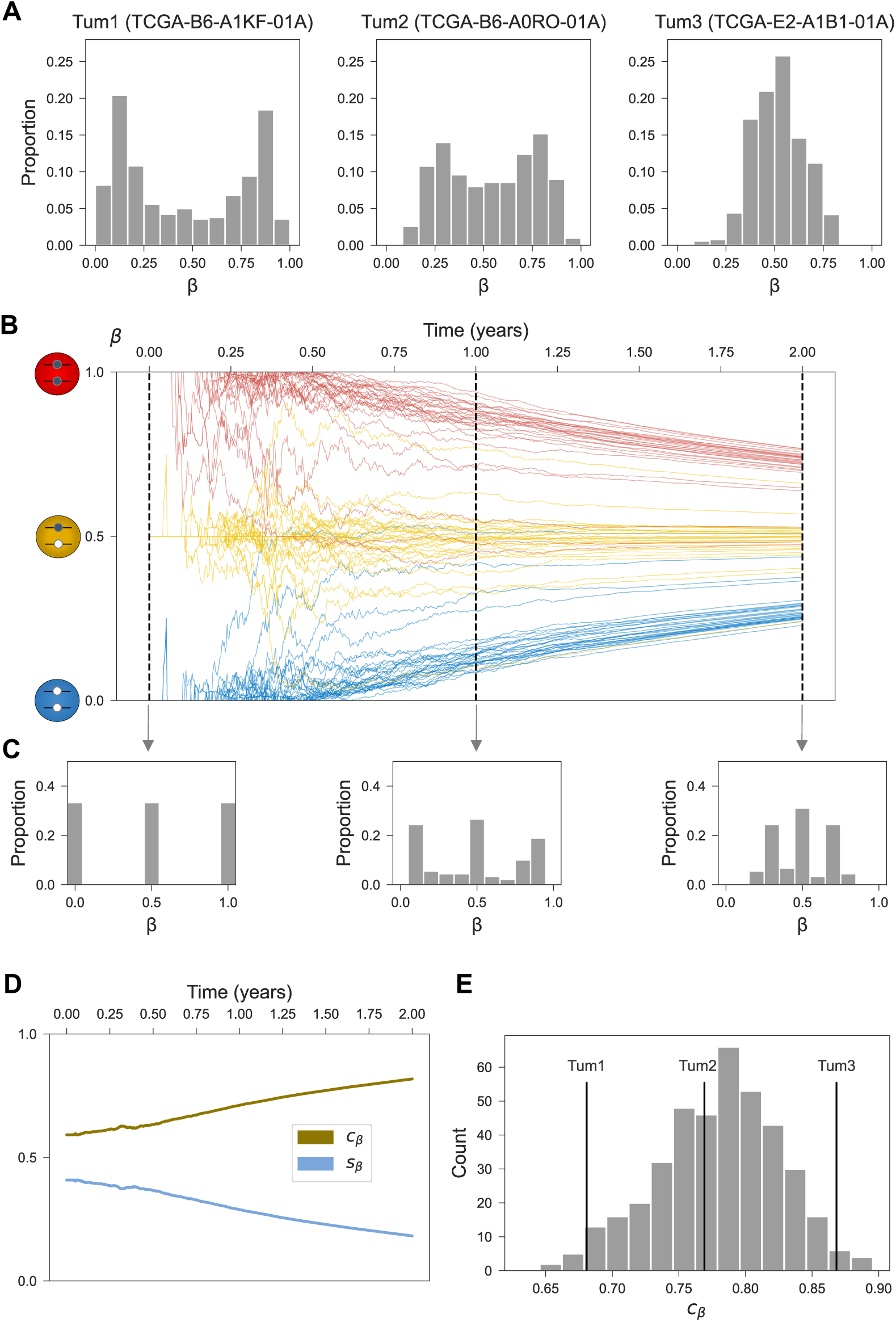
The distribution *β*-values across the clock set encodes tumor age. **(A)** The empirical *β*-value distributions for the clock set fCpGs (N=500) are shown for three select tumors in the TCGA cohort. **(B)** Simulated trajectories for an ensemble of fCpG sites (N=90), starting in the unmethylated, hemi-methylated, and methylated initial configurations, respectively (n=30 each). The thick lines represent the average *β*-value trajectories for each subset. Simulation parameters as detailed in Figure 1. **(C)** Cross-sectional *β*-value distributions for the simulated clock set in panel B, shown after 0, 1, and 2 years of growth. **(D)** Standard deviation (*s_β_*) of *β*-values and epigenetic clock index (*c_β_* = 1 - *s_β_*) over 2 years of growth for the simulated clock set in panel B. **(E)** The distribution of epigenetic clock index values (*c_β_*) across invasive ductal carcinomas in TCGA (N=400); the three tumors from (A) are labeled.

By considering the histograms of *β*-value distributions at different mitotic ages, we can track the evolution of the three “peaks” corresponding to the subsets of initially unmethylated, hemi-methylated, and methylated clock sites (**Figure 3C**). As the tumor’s mitotic age increases, the left peak of the histogram (consisting of originally unmethylated clock sites) starts moving to the right, whereas the right peak (originally methylated sites) moves to the left; the middle peak (originally hemi-methylated sites) remains stationary. By measuring the extent to which the three peaks have converged to the stationary value of *β* = 0.5, we can thus estimate the mitotic age of individual tumors.

Concretely, we used the standard deviation of the *β*-values, denoted by *s_β_*, to quantify the relationship between mitotic tumor age and the evolving clock set profile (**Figure 3C**). Because *s_β_* is highest at time 0, when the *β*-value distribution exhibits three sharp peaks, and then monotonically decreases over time (**Figure 3D**), we introduced the epigenetic clock index *c_β_* = 1 − *s_β_* as a proxy measure of mitotic tumor age (**Figure 3E**).

In the next two sections, we characterize the relationship between a tumor’s mitotic age, as quantified by the epigenetic clock index *c_β_*, and its evolutionary-ecological context as determined by its intrinsic growth potential and external pressures from the microenvironment. Because there is a pronounced difference in *c_β_* between ductal carcinomas (median, 0.79) and lobular carcinomas (median, 0.82; P=2×10^−16^, Wilcoxon rank-sum test), we restricted subsequent analyses to the more common ductal carcinomas to avoid unnecessary confounding.

### Younger tumors have more aggressive phenotypes

As a breast tumor grows, its likelihood of detection on the basis of imaging or symptoms increases. Because fast growing tumors are expected to reach a detectable size sooner than slow growing ones, we hypothesized that younger tumor mitotic age would correlate with established markers of tumor aggressiveness. To test this hypothesis, we correlated the epigenetic clock index with several established features of tumor aggressiveness, including molecular subtype,^22,23^ genomic instability,^24,25^ grade,^26^ and size.^27^

There was a clear relationship between mitotic age and molecular subtype: luminal A tumors, which have a more favorable prognosis, were older than luminal B and basal tumors (**Figures 4A-B; Suppl. Table S1**). Similarly, there was a strong correlation between genomic instability and younger tumor age (**Figures 4C-D**), and younger tumors were of higher histopathologic grade (**Figure 4E**). In contrast, there was only a weak relationship between mitotic age and summary stage (**Figure 4F**)

**Figure 4.**
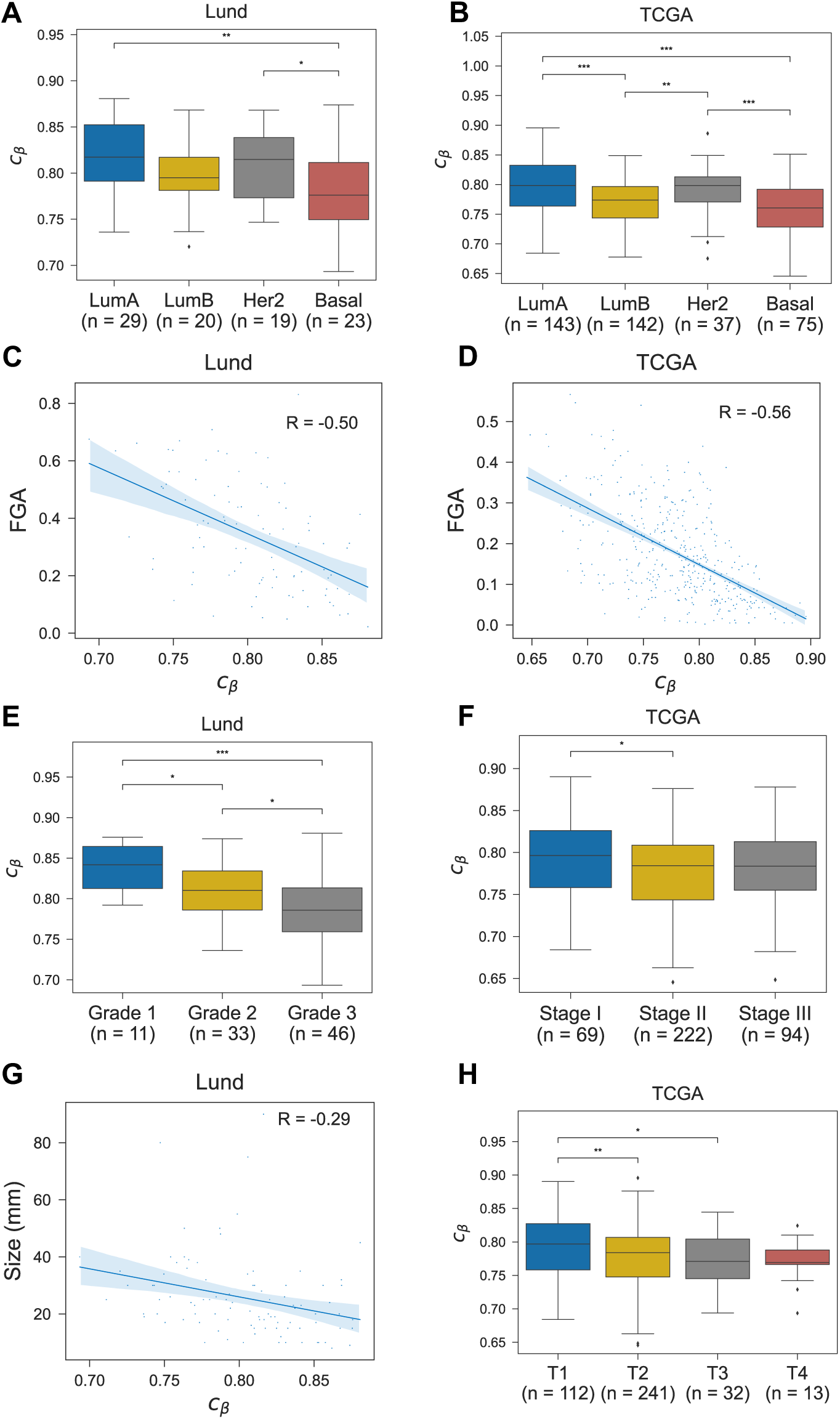
Epigenetic clock index vs. clinicopathological variables. The distribution of epigenetic clock index values (*c_β_*) among invasive ductal carcinomas in the Lund and TCGA cohorts, by **(A)** tumor grade; **(B)** tumor stage; **(C)-(D)** molecular subtype as predicted by the PAM50 algorithm; **(E)** tumor size; **(F)** T-stage; and **(G)-(H)** fraction of genome altered (FGA) by copy number alterations. Pairwise comparisons of medians in panels A, B, E, F, and H were performed using a two-sided Wilcoxon rank-sum test (*P< 0.05, **P< 0.01, ***P< 0.001). In panels C, D, and G, regression lines and bootstrapped 95% confidence intervals are shown; Pearson correlations (R) are indicated.

Another prognostic factor in breast cancer is tumor size, with larger lesions having worse outcomes. We found that smaller tumors were of older mitotic age compared to larger tumors (**Figures 4G-H**), presumably because slow growing tumors spend more time at the smaller end of the detectable size range, and are, therefore, more likely to be detected at a smaller size.

Finally, the relationship between tumor mitotic age and patient age at diagnosis was inconclusive, with a weak negative correlation in TCGA (R=-0.18) and no correlation in the Lund cohort (R=-0.06; **Suppl. Table S1**). This is consistent with the notion that the fCpG clock measures the age of the tumor—starting with the most recent common ancestor cell—and not the age of the patient.

### Identifying modulators of mitotic tumor age

The time it takes for a tumor to grow from a single cell to a detectable mass depends on its effective growth rate, that is the difference between cell proliferation and cell death (**Figure 5A**). Cell proliferation primarily reflects the tumor’s intrinsic growth potential and aggressiveness, whereas cell death is often the result of extrinsic selective pressures applied by the tumor microenvironment, such as immune surveillance and resource constraints due to limited vascularization.^28,29^

**Figure 5.**
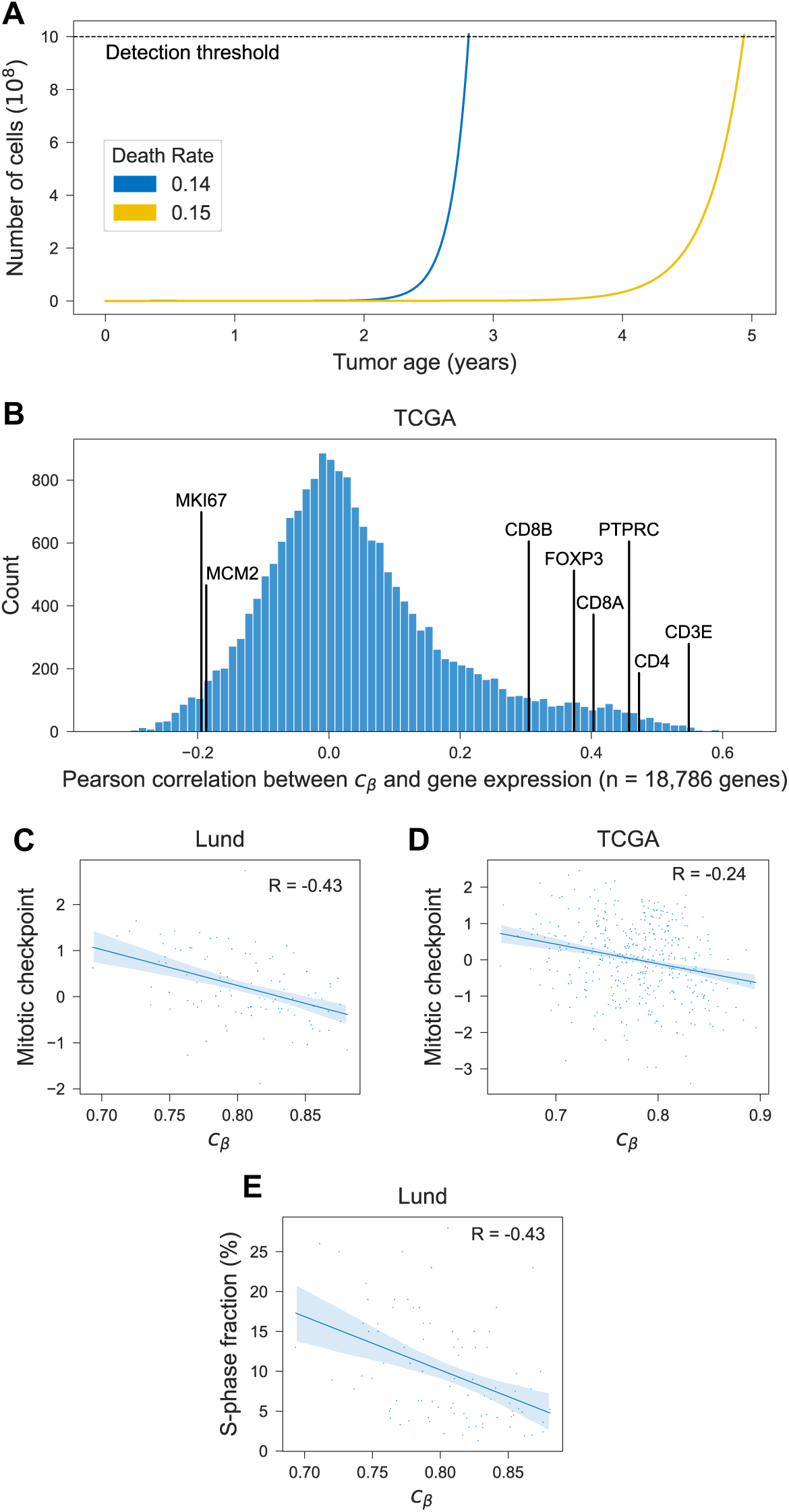
Mitotic tumor age vs. measures of proliferation. **(A)** Simulation of tumor size as a function of tumor age. Both tumors have the same proliferation rate (*α* = 0.16 divisions/day), but different death rates: *λ* = 0.14 1/day (blue) vs. *λ* =0.15 1/day (yellow). **(B)** Pearson’s correlation between epigenetic clock index *c_β_* and expression of protein-coding genes (TCGA cohort); correlation with select genes as indicated. **(C)-(D)** Correlation of *c_β_* with average expression of genes involved in M-phase and mitotic checkpoint regulation. **(E)** Correlation of *c_β_* with the fraction of cells in S-phase, as measured by flow cytometry (Lund cohort). Regression lines shown with bootstrapped 95% confidence intervals and Pearson correlation (R).

To explore putative modulators of effective tumor growth and mitotic age at diagnosis, we performed genome-wide correlation analyses of the epigenetic clock index *c_β_* against gene expression. As predicted, mitotically younger tumors exhibited increased expression of proliferation-related genes such as Ki67 and MCM2 (**Figure 5B, Suppl. Table S1**). The signal was further augmented when considering the average expression across a set of genes involved in M-phase and mitotic checkpoint regulation (**Figure 5C-D**) and the fraction of cells in S-phase (**Figure 5E**).

Next, we examined the microenvironment’s ability to decrease the effective growth rate of a tumor through increased cell death. As hypothesized, the expression of immune cell markers such as CD3, CD4, CD8 and FOX3 was elevated in mitotically older tumors (**Figure 5B; Suppl. Table S1**). This suggests that tumors which are subject to immune surveillance—e.g., through neo-antigen directed immune control by CD8+ T-cells—have a lower effective growth rate and, thus, reach a detectable size at an older mitotic age, as compared to tumors that successfully evade immune control and thus reach a detectable size at a younger mitotic age.

To perform a systematic analysis of mitotic tumor age modulation, we performed a genome-wide gene set enrichment analysis (GSEA) (**Figure 6A**). Consistent with the univariate gene expression analyses, mitotically younger tumors were enriched for pathways related to proliferation and cell cycle control. Conversely, mitotically older tumors were enriched for immune pathways and immune-related signaling pathways, again supporting the notion of effective immune control in older, slower growing lesions.

**Figure 6.**
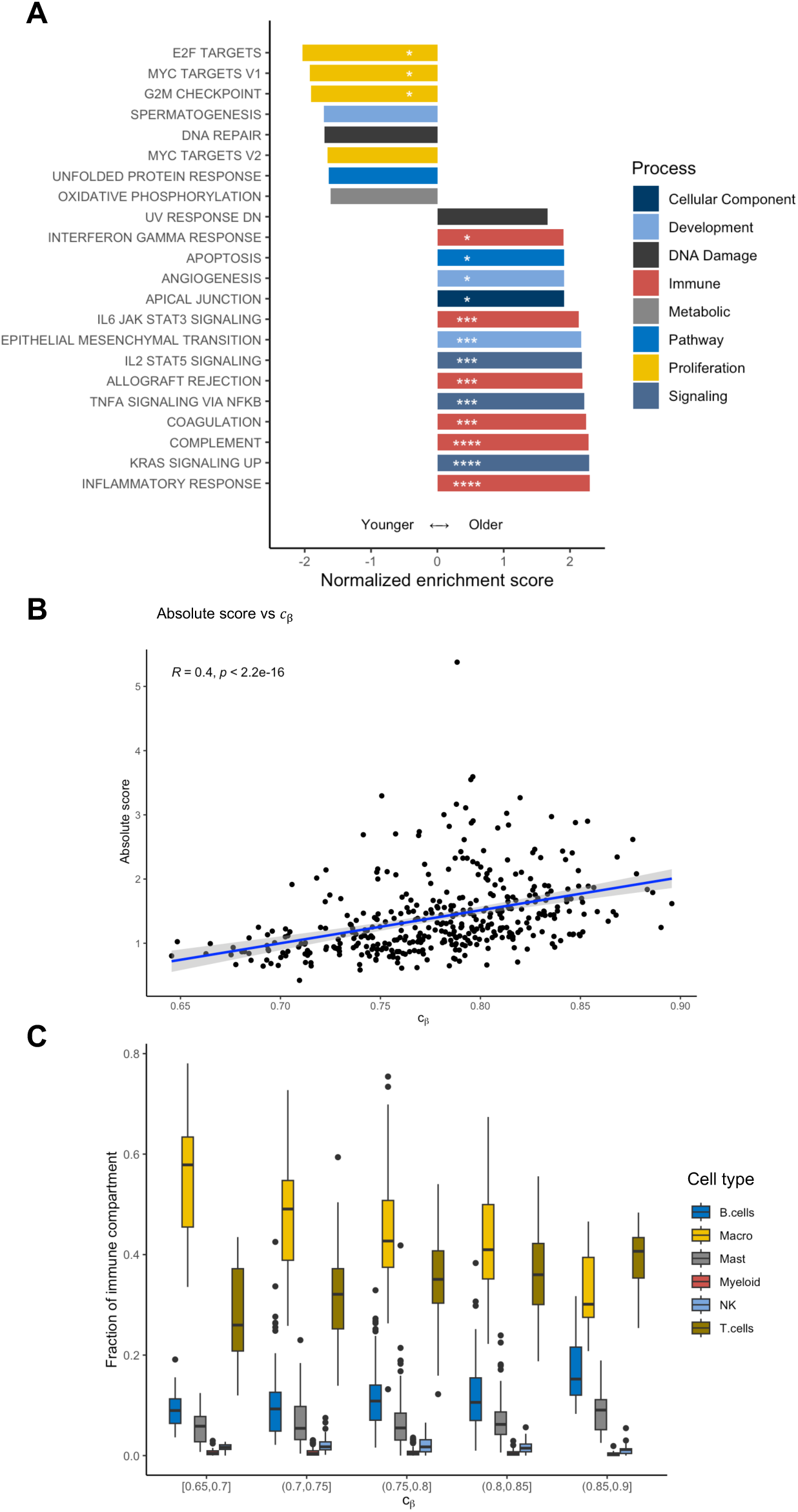
Pathway enrichment and immune decomposition analyses. **(A)** A gene set enrichment analysis (GSEA) was performed for the epigenetic clock index *c_β_*. Pathways with a positive (negative) enrichment score are enriched in mitotically older (younger) tumors. Only pathways with a false discovery rate (FDR) below 0.1 are shown; *FDR<.05; **FDR<.01; ***FDR<.001; ***FDR<.0001. **(B)** Epigenetic clock index vs. the extent of immune infiltration (absolute immune score) as estimated by CIBERSORTx. **(C)** The immune compartment of each tumor was decomposed using CIBERSORTx; the compartment fractions are shown for tumors of similar mitotic age (epigenetic clock index *c_β_*).

For a more in-depth analysis of the immune infiltrate, we used the CIBERSORTx algorithm^30^ to estimate the extent and composition of the immune compartment. As expected, the extent of the immune compartment increased with mitotic tumor age (**Figure 6B**). When decomposing each tumor’s immune compartment into the major cell types, we found an increase in the fraction of T-cells in mitotically older tumors (**Figure 6C, Suppl. Table S2**), again suggestive of T-cell mediated immune surveillance.

### Analysis of paired tumor samples validates epigenetic clock

Multiple tumor samples from the same patient provide a unique opportunity to assess the internal validity of the epigenetic clock. Indeed, paired samples should be epigenetically more related—via their most recent common ancestor cell—than samples from different patients. In a cohort of 8 women with multi-focal breast cancer,^31^ we found that the within-patient correlations of the clock set fCpG sites were higher (median, 0.72) than the between-patient correlations (median, 0.10; P=3×10^−6^, Wilcoxon rank-sum test; **Figure 7A**). The same held true for a cohort of 18 patients with paired primary tumors and lymph node metastases^32^ (median, 0.82 vs. 0.14, P=5×10^−13^; **Figure 7B**) and a subset of 22 patients with paired primary tumors and metastases (including lymph node and distant metastases) from the AURORA US Metastasis Project^33^ (median, 0.73 vs. 0.08, P=2×10^−26^; **Suppl. Figure S4**).

**Figure 7.**
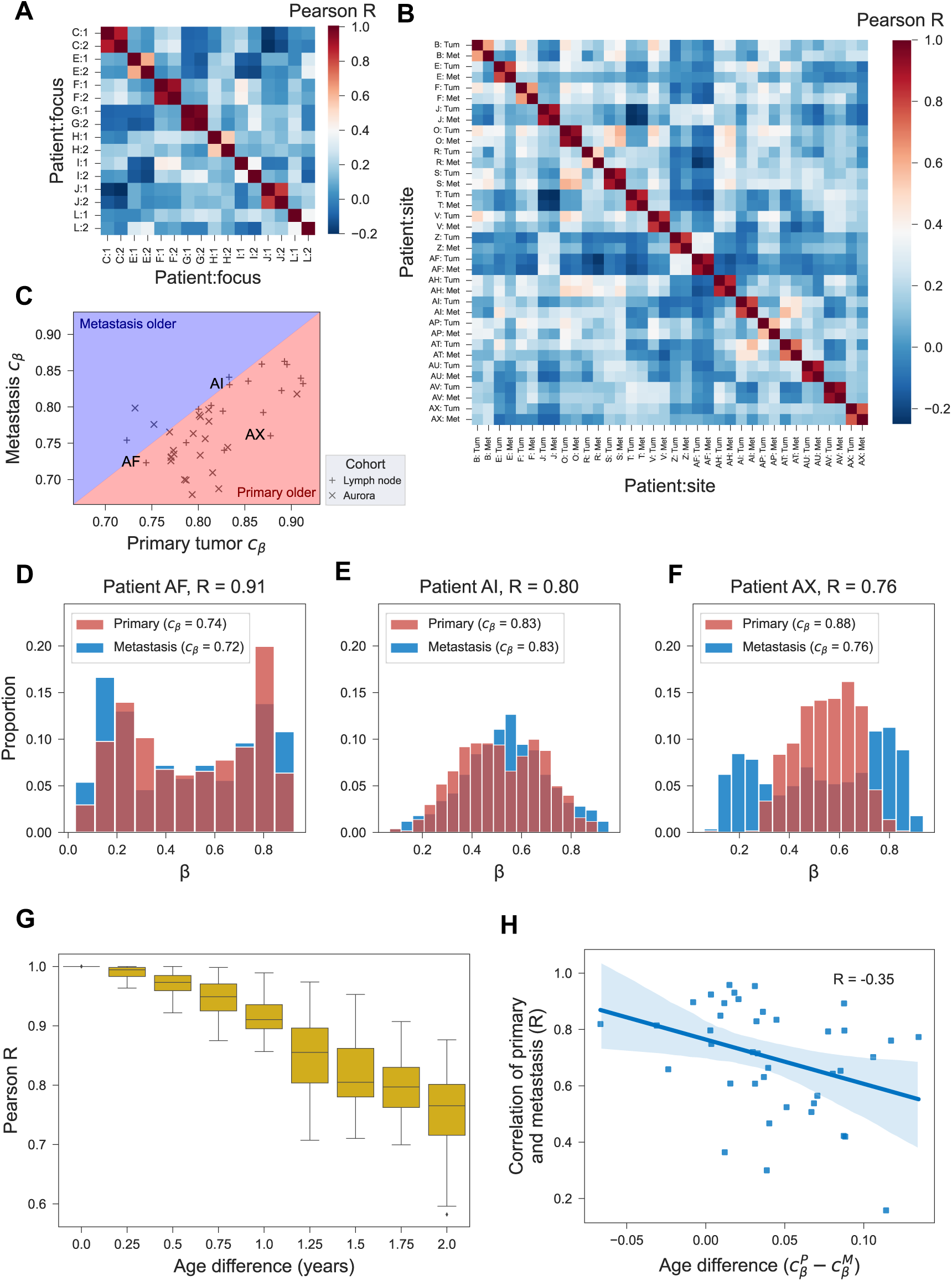
Paired tumor samples. **(A)** In a cohort of 8 patients with multifocal breast cancer, the *β*-values of the 500 fCpG sites of the epigenetic clock are correlated within (two foci per patient) and between patients. **(B)** In a cohort of 18 patients with paired primary tumor and lymph node metastasis samples, the *β*-values of the 500 fCpG sites are correlated within and between patients. **(C)** For the 18 patients from panel B and 22 patients from the AURORA US Metastasis Project, the epigenetic clock index of the primary tumor is plotted against the index of the lymph node metastasis. Three patients (AF, AI, and AX) from the lymph node cohort are labeled for the purposes of the following 3 panels. **(D-F)** For the 3 labeled patients from panel C, the *β*-value distributions of fCpGs are shown both for the primary tumor and the metastasis. **(G)** Monte Carlo simulation of an ensemble of 90 fCpG sites, 30 each starting in the unmethylated, hemi-methylated, and methylated initial configurations; every 3 months, a cell was randomly picked to representing the metastasis seeding cell, and the Pearson correlation between the *β*-value distribution of that cell and that of the entire tumor was calculated. Distributions at each time point represent the results from 30 independent simulations. Simulation parameters are detailed in Figure 1. **(H)** For the 18 patients from panel B and 22 patients from the AURORA US Metastasis Project, the epigenetic relatedness of primary and lymph node metastasis (Pearson’s R for the 500 fCpG sites in the clock set) is compared to the difference in epigenetic clock index, as a proxy for the difference in mitotic age between the two samples.

The two cohorts of patients with paired primary and metastasis samples^32,33^ allowed us to test two additional properties of the fCpG clock. First, assuming that each metastasis is seeded by a single cell from the primary tumor, synchronous metastases should be younger than their matched primaries. Indeed, the epigenetic clock measures the age of the metastasis relative to the seeding event, which occurred after initiation of the primary tumor. Consistent with this prediction, in 36/40 patients, we found the metastases to have a lower epigenetic clock index compared to their matched primaries (**Figure 7C**). This provides direct support for our interpretation of the epigenetic clock index as a proxy measure for mitotic age.

Second, the timing of metastatic dissemination relative to the primary tumor’s age is expected to impact the epigenetic similarity of the two samples: if the metastasis is seeded early during primary tumor growth (i.e., similar *c_β_* values), the *β*-values of the two samples are expected to be closely related (**Figures 7D-E**) because the metastasis seeding cell came from a mostly homogenous population; conversely, if the metastasis is seeded late (i.e., different *c_β_* values), the *β*-values are expected to differ more substantially (**Figure 7F**) because the seeding cell came from a heterogenous population. Corroborating this hypothesis, and consistent with a corresponding simulation of metastatic seeding based on the oscillator model (**Figure 7G**), we found a negative correlation between mitotic age difference and *β*-value similarity (**Figure 7H**).

### Quantifying mitotic tumor age

So far, we have used the epigenetic clock index *c_β_* as a correlate of mitotic tumor age. To derive quantitative estimates of each tumor’s mitotic and calendar ages, we proceeded as follows (see **Methods** for details). First, we invoked the mathematical oscillator model (**Figure 1A**) to relate mitotic tumor age to the measured *β*-values of fCpG sites in the clock set. Next, we decomposed each tumor’s empirical fCpG *β*-value distribution into three groups (**Figure 8A**): originally unmethylated fCpGs (left peak in the histogram), originally hemi-methylated fCpGs (middle peak), and originally methylated fCpGs (right peak). Finally, we combined the peak location in each group with the oscillator model to infer the estimated mitotic age of the tumor (**Figure 8B**).

**Figure 8.**
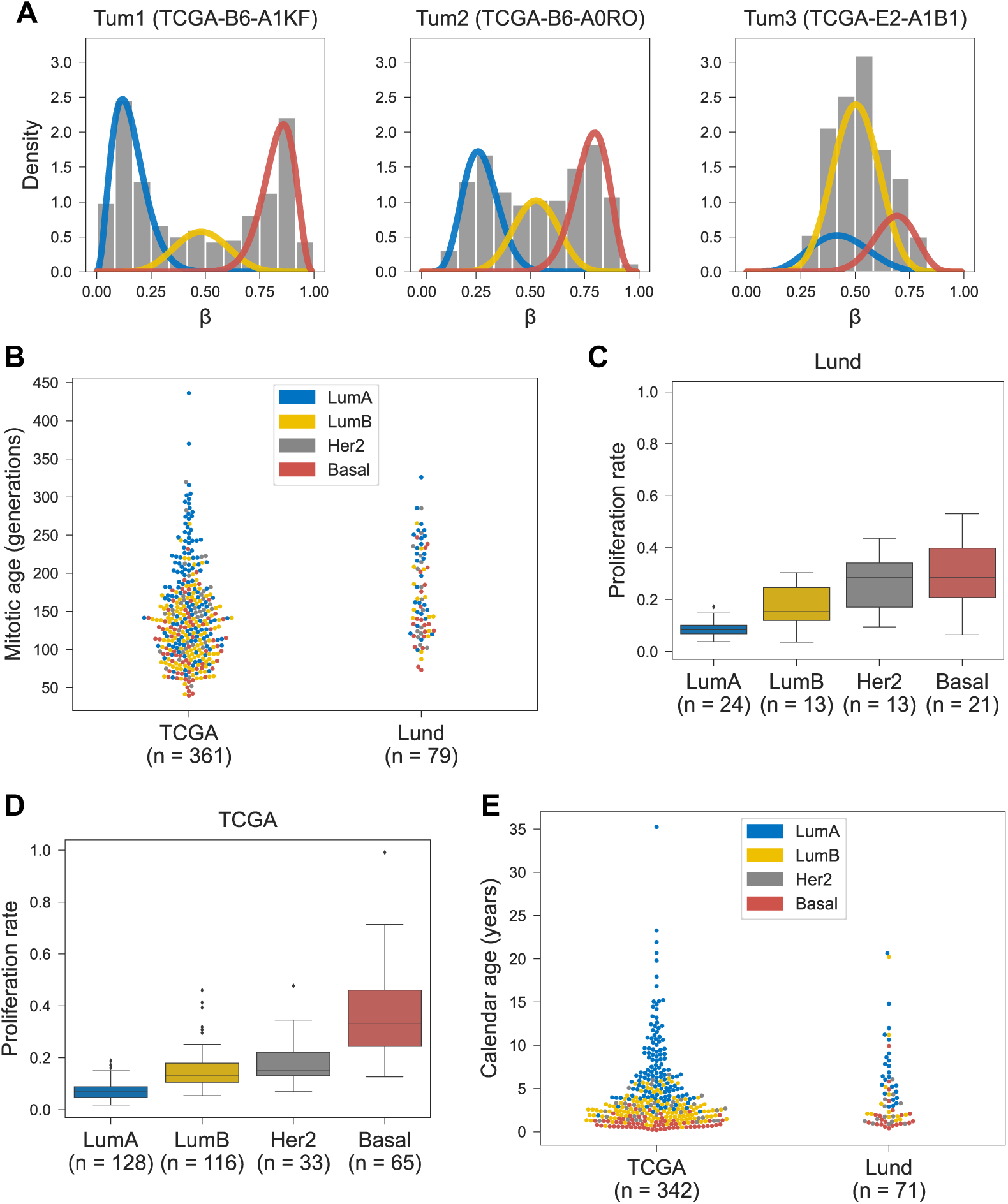
Estimating tumor age. **(A)** The empirical *β*-value distributions of the 500 fCpG sites in the clock set are decomposed into three peaks corresponding to initially unmethylated (blue), hemi-methylated (yellow), and fully methylated (red) sites. **(B)** Mitotic tumor age (the number of generations) is shown for the TCGA and Lund cohorts and colored by molecular subtype. These estimates are based on the peak locations (panel A) and a (de-)methylation rate of µ = 2 ⋅ 10^−3^, per cell division and per allele, which yields a mean tumor age of 3 calendar years (see panel E). **(C)** For the Lund cohort, tumor-specific proliferation rates are estimated using the measured fraction of cells in S-phase and shown by molecular subtype. **(D)** For the TCGA cohort, proliferation rates are estimated using S-phase fractions predicted based on gene expression, using a model trained on the Lund cohort. **(E)** Calendar age of tumors in the TCGA and Lund cohorts, colored by molecular subtype.

Finally, we combined tumor-specific estimates of mitotic age (**Figure 8B**) and proliferation rate (**Figures 8C-D**) to derive tumor-specific estimates of calendar age. Anchoring the median tumor age at a consensus estimate of 3 years (see **Methods**), the distribution of calendar ages across the TCGA and Lund cohorts ranged from 0.2 to 35.2 years, with an interquartile range of 1.5 to 5.6 years (**Figure 8E**). There were notable differences in median tumor calendar ages by molecular subtype, ranging from 1.0 years in basal cancers to 6.5 years in Luminal A cancers (**Suppl. Table S3).**

### Adjusting for tumor purity

Bulk samples contain a mixture of tumor and stroma. Because the epigenetic clock index exhibited correlations with tumor purity as measured by the consensus purity estimate^34^ (CPE; R=-0.67; **Suppl. Figure S1A**), we restricted our analyses to samples of high tumor purity (CPE≥0.6). Nevertheless, we cannot rule out that the observed variability in *β*-value distributions among the selected fCpG sites—which are used to estimate mitotic tumor age—were at least partially driven by the methylation patterns of admixed non-epithelial cells. If this is the case, then, e.g., the immune pathway enrichment of older tumors (**Figure 6A**) may be confounded by the presence of non-epithelial cells that alter the measured *β*-value distribution.

To adjust for possible confounding by tumor purity, we derived purity-adjusted *β*-values for the tumor cells by modeling the measured methylation as a mixture of tumor and stroma methylation, see **Methods** for details. The resulting purity-adjusted epigenetic clock index 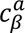 exhibited a lower correlation with tumor purity (R=-0.22; **Suppl. Figure S1B**) and was lower than the unadjusted epigenetic clock index *c_β_* (**Suppl. Figure S1C**).

When replacing the unadjusted epigenetic clock index with the purity-adjusted version, the strength of correlations between markers of tumor aggressiveness and younger mitotic age remained unaltered (**Suppl. Table S1, Suppl. Figure S2, Suppl. Figure S3**). Individual immune genes and the extent of immune infiltration remained associated with older mitotic age, although the correlations were attenuated (**Suppl. Table S1, Suppl. Table S2**). While the immune pathways were no longer enriched in older tumors (**Suppl. Figure S3**), there was still a positive correlation between the fraction of T cells and mitotic age (**Suppl Table S2**).

## DISCUSSION

In this study, we developed an epigenetic clock to measure the age of newly diagnosed breast cancers. Measuring epigenetic entropy among neutrally fluctuating CpG (fCpG) sites, the clock tracks mitotic tissue age on a time scale of years and, thus, provides higher temporal resolution compared to previous tissue clocks. Based on standard methylation arrays, it has the potential to be a novel marker of aggressiveness and prognosis in early-stage breast cancer.

Once a patient is diagnosed with breast cancer, the tumor’s mitotic age encodes valuable prognostic information. Intuitively, a slow-growing tumor that takes a long time to reach the threshold of detection is more likely to have a good prognosis compared to a fast-growing tumor that quickly expands into a detectable mass. Our analyses corroborate this hypothesis by revealing that mitotically younger tumors were enriched for features of tumor aggressiveness and predictors of poor outcome, including genomic instability, higher grade, and basal molecular subtype.^24,25^ This property of the epigenetic clock is quite remarkable given that its constituent fCpG sites were selected from non-functional regions of the genome, on the basis of simple statistical properties of their *β*-value distributions.

Beyond prognostication, the clock holds promise in risk-stratified screening approaches. The efficacy of breast cancer screening critically depends on the sojourn time, that is the time window during which the tumor is asymptomatic but mammographically detectable. If the sojourn time is short, early detection is unlikely even under frequent screening; if it is long, some cancers will be overdiagnosed.^35^ Sojourn time estimates are usually obtained by fitting natural history models to population data, yielding indirect, population-averaged estimates. Our approach, in contrast, allows for direct and individual-level characterization of tumor age, which provides a lower bound for the sojourn time. Assuming an overall median time to detection of 3 years in our cohort, the time to detection in luminal A cancers (6.5 years) was substantially longer compared to that in Luminal B (2.4 years) and basal (1.0 years) cancers. These estimates are consistent with the observation that interval cancers are enriched for more aggressive subtypes compared to screen-detected cancers,^36^ and highlight opportunities for data-driven personalization of screening schedules.

The epigenetic clock also provides an opportunity to quantify the evolutionary-ecological pressures that shape the temporal landscape of breast cancers. Indeed, because most tumors are of comparable size at the time of diagnosis, mitotic age is related to the effective growth rate: tumors that reach the detection threshold at a younger age have a higher effective growth rate compared to tumors that reach the threshold at a higher mitotic age. Our analyses characterized the effective growth rate of breast cancers as a competition between tumor-intrinsic growth potential (e.g., proliferation) and microenvironmental pressures (e.g., surveillance by immune cells).^37^ According to this model, highly proliferative tumors that successfully evade the immune system are detected at a younger age compared to less proliferative lesions subject to continuous immune control.

Our study has several limitations. First, because tumor age is not observable in practice, a direct validation of the clock is not possible. Nevertheless, we note that the clock correctly classified the age ordering of primary tumors and metastases in 36 of 40 patients. Second, the epigenetic clock index was correlated with sample purity, which suggests the latter may be a confounder in our analyses. To mitigate this risk of bias we systematically repeated all analyses using purity-adjusted methylation values; while some of the associations were attenuated, the overall qualitative conclusions remained unchanged. To address the potential confounding of age estimates by tumor purity, single cell methylation data is needed. Third, estimation of mitotic tumor age was based on a simple mathematical model of (de-)methylation dynamics. In future work, this approximation can be refined using more sophisticated simulation-based models that account for the underlying population dynamics, including cell proliferation and death, and possibly selection.

How long a newly diagnosed breast cancer has been growing is generally considered a known unknown. Here we revisited this assumption and developed a new way to infer tumor age using standard methylation arrays. While developed specifically for breast cancer, the approach can be generalized to any cancer type and, as such, provides a scalable technology to characterize the temporal landscape of oncology.

## METHODS

### TCGA Cohort

Of the 1,085 invasive breast cancers from female patients in The Cancer Genome Atlas (TCGA),^38^ 774 had available methylation array data (Infinium HumanMethylation450 BeadChip, Illumina, San Diego, CA, USA). After excluding 138 tumors of low tumor content (consensus purity estimate^34^ [CPE] <0.6) and one sample each from two patients with two primary samples, the remaining 634 tumors were used to select the ensemble of 500 fCpG sites. Finally, after excluding 10 tumors with ≥5% missing clock set fCpG measurements and 100 tumors with a histology code other than *infiltrating duct carcinoma*, the analytic cohort consisted of 400 tumors. The following variables were retrieved: patient age at diagnosis; tumor histology; T stage (subsetted to T1, T2, T3, and T4); summary stage (subsetted to stages I, II, or III). For all 400 patients with invasive ductal carcinoma, gene expression quantification (RNA-seq) and copy number segment data were available as well; when >1 measurement was available, one was selected at random. All clinical and sequencing data were retrieved from the Genomic Data Commons (GDC; https://gdc.cancer.gov) using the R package *TCGAbiolinks* (version 2.25.3).

### Lund cohort

We retrieved publicly available methylation array data (Infinium HumanMethylation450 BeadChip) from 181 primary breast cancers in the Southern Sweden Breast Cancer Group tissue bank at the Department of Oncology and Pathology, Skåne University Hospital (Lund, Sweden) and the Department of Pathology, Landspitali University Hospital (Reykjavik, Iceland).^20^ The data were obtained through the Gene Expression Omnibus (GSE75067). Because calculation of the purity metric CPE requires gene expression, somatic copy-number, and immunohistochemistry in addition to methylation data, we instead assessed tumor purity using the leukocyte unmethylation percentage (LUMP) value. A tumor’s LUMP value is calculated as the average *β*-value among 44 specific CpG sites, divided by 0.85; we found the LUMP value to be strongly correlated with CPE (R=0.86). After excluding samples of low purity (LUMP <0.6; n=35), the remaining 146 samples all had ≤5% missing clock set fCpG measurements. After exclusion of non-ductal histology (n=48) we ended up with an analytic cohort of n=98. The following variables were retrieved: patient age at diagnosis; tumor grade; tumor size; molecular subtype (PAM50); fraction of genome altered (FGA); expression of a mitotic checkpoint gene module;^39^ fraction of cells in S-phase (flow cytometry).

### Normal breast tissue cohort

We obtained publicly available methylation array data (Infinium HumanMethylation450 BeadChip) from 100 normal breast tissue samples in the Susan G. Komen Tissue Bank ^17,40^ (GSE88883). We excluded samples of low purity (LUMP<0.6), resulting in a cohort of 79 normal breast tissue samples used for identifying fCpG sites.

### Multiple sample cohorts

We retrieved publicly available methylation array data (Infinium Human-Methylation450 BeadChip) from four cohorts with paired tumor samples. The first cohort consisted of 8 breast cancer patients with multiple primary samples (GSE106360).^21^ Only samples from the 5 patients (2 patients with 5 samples each; 3 patients with 3 samples each) who had not received neoadjuvant therapy were used. Because LUMP values were highly variable, we did not apply any purity filtering. The second cohort consisted of 10 patients diagnosed with multi-focal breast cancer (GSE39451).^31^ For each patient, methylation array data from 2 foci were available, and we only included the 8 patients where both samples were of sufficient purity (LUMP≥0.6). The third cohort consisted of paired primary and lymph node metastasis samples from 44 patients (GSE58999).^32^ Only patients where both samples were of sufficient purity (LUMP≥0.6) were included (n=18). The fourth cohort, from the AURORA US Metastasis Project, consisted of primary and metastasis samples taken from 55 patients with metastatic breast cancer. In our analysis, we included only patients for whom at least one primary and one metastasis sample of sufficient purity (LUMP≥0.6) were available (n=22) (GSE212370).^33^ Only patients with at least one primary and one metastasis sample of sufficient purity (LUMP≥0.6) were included (n=22). When more than one primary or metastasis sample was available, the one with the highest LUMP value was selected.

### Selection of fluctuating CpG (fCpG) sites

First, we identified CpG sites on the HumanMethylation450 BeadChip that correspond to functional regions of the genome. To this end, we identified sites that were associated with regulatory features or genes in one or both of the official annotation files of the Infinium HumanMethylation450 and MethylationEPIC bead chip arrays (https://support.illumina.com). After exclusion of such functional sites, a total of 86,099 CpG sites without functional annotation remained. Next, we sought to identify CpG sites with balanced methylation and demethylation rates, defined as having an average methylation content (*β*-value) between 0.4 and 0.6 in both the TCGA cohort (N=634) and the normal breast tissue cohort (N=79). CpG sites with ≥20 missing values in either cohort were excluded from this selection process. In the last step, we ranked all balanced CpG sites by their *β*-value variance among tumors in the TCGA cohort and selected the 500 most variable fCpGs to define the clock set 𝒞. Based on the clock set, each tumor was assigned an epigenetic clock index *c_β_* = 1 − *s_β_* where *s_β_* is the standard deviation of the *β* values in the clock set.

### Gene expression analyses

For tumors in TCGA, relative gene expression levels were taken as the mean-centered, log2(x+1) transformation of the reported transcript per million (TPM) intensities. Among the 60,616 RNA transcripts recorded in TCGA, only those classified as protein-coding genes by the HUGO Gene Nomenclature Committee^41^ were included in subsequent analyses (n=18,910). Expression of MCM was calculated as the average relative expression in the 6-gene family MCM2-7, and expression of a *mitotic checkpoint* gene module^39^ was calculated as the average relative expression of the genes included in the module. Molecular subtyping was based on the PAM50 algorithm^42^ as implemented in R package *Genefu*.^43^ For tumors in the Lund cohort, identical gene expression and molecular subtyping analyses had previously been reported,^39^ thus enabling a direct comparison between tumors in the TCGA and Lund cohorts.

### Pathway enrichment analyses

For the TCGA cohort, we performed a gene set enrichment analysis (GSEA) using the software package *GSEA*^44,45^ to identify *Hallmark* gene sets that are correlated with the epigenetic clock index *c_β_*. The analysis was performed using the Pearson correlation to rank individual genes; phenotype-permutation-based *P* values and false-discovery rate (FDR) *Q* values were computed using 1,000 permutations. All other inputs were kept at their defaults.

### CIBERSORTx

To assess the immune cell composition within the tumor microenvironment, we employed CIBERSORTx using the LM22 signature matrix and batch correction.^30^ Briefly, RNA-seq data from the TCGA tumor samples were uploaded to the CIBERSORTx web portal, where gene expression profiles were deconvoluted to estimate the absolute scores for 22 distinct immune cell types. The analysis was performed with the default parameters, including 100 permutations for statistical significance assessment. For reporting of results the 22 distinct cell types were then collapsed into six mutually exclusive categories: B cells, macrophages, mast cells, myeloid cells, natural killer (NK) cells, and T cells.

### Copy-number analyses

For tumors in TCGA, copy-number (CN) data consisted of specified chromosomal regions of equal CN, the 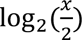-transformed CN, and the number of probes. We converted these values to absolute copy numbers and determined each segment to have either a copy number gain (segment mean ≥ 2.5), a copy number loss (segment mean ≤1.5), or no change (1.5 < segment mean < 2.5). The fraction of the genome altered by copy number gains and losses were each calculated for every tumor by dividing the number of probes affected by gains and losses, respectively, by the total number of probes. The total fraction of the genome altered by copy number alterations (FGA) was then calculated as the sum of these two values. For tumors in the Lund cohort, the same approach had previously been used to compute FGA,^2^ thus enabling direct comparison between tumors in the TCGA and Lund cohorts.

### In silico model of tumor growth and fCpG dynamics

To simulate the dynamics of fCpG sites in a growing tumor, we used a discrete-time birth-death process. Starting with a single founding tumor cell, the population is updated in time intervals of one day, at which time each cell either divides, dies, or remains unchanged with probabilities *α*, *λ*, and 1 − *α* − *λ*, respectively. Upon cell division, each allele in each cell changes its methylation state with probability *μ*. We tracked an ensemble of 90 fCpG sites, assuming independent (de-)methylation dynamics. Unless otherwise specified, the following parameters were used: *α* = 0.17 (the estimated mean proliferation rate in the TCGA-Lund combined cohort, see below for details), *λ* = 0.15 (to reach a population of 10^9^ cells in 3 years, or, (1 + *α* − *λ*)^3⋅365^ ≈ 10^9^), and µ = 0.002 (the estimated flip rate in the combined cohort, see below for details).

### Tumor specific proliferation rates

For tumors in the Lund cohort, tumor specific proliferation rates *α_i_* were estimated based on the reported fraction *f_i_* of cells in S-phase as *α_i_* = *f_i_*/*T_s_*, where *T_s_* is the average time spent in S-phase (see Supplementary Methods for details). We assumed *T_s_* to equal 12.7 hours, based on an average across five cancer cell lines.^46^ Because *f_i_* is not reported in the TCGA cohort, we used the Lund data to develop a predictive model of S-phase fraction using an elastic net model. As candidate predictors, we included FGA, LUMP value, and average gene expression levels within each of the following gene modules:^39^ mitotic checkpoint (see above), immune response, stroma, mitotic progression, early response, steroid response, basal, and lipid. The model was fit to the Lund cohort tumors using cross-validation for hyperparameter optimization, and then applied to TCGA tumors to predict tumor-specific S-phase fractions *f_i_* and proliferation rates *α_i_*.

### Tumor age estimation

To estimate tumor mitotic and calendar ages from the empirical *β*-value distributions, we proceeded in two steps. In the first step, we decomposed each tumor’s empirical *β*-value distribution into three groups, or “peaks”, of fCpG sites: the originally unmethylated fCpG sites (left peak), the originally hemi-methylated fCpG sites (middle peak), and the originally methylated fCpG sites (right peak). We achieved this by fitting a mixture model of three Beta distributions to the *β*-values of the 500 fCpG sites in the clock set using the R package *BetaModels* (version 0.5.2). To improve convergence of this method, sites with extreme *β*-values (*β* > 0.98 or *β* < 0.02) were removed before fitting the mixture model (a total of 69 and 442 sites were thus removed in the TCGA and Lund cohorts). In preparation of the next step, we determined the mode of each Beta component in the mixture as the location of the corresponding peak. At this point we excluded tumors with a middle peak location outside the interval [0.4, 0.6] because this suggests a bias in the (de-)methylation rates and thus violates a basic assumption of the fCpG dynamics in the clock set (44 and 12 tumors were excluded in the TCGA and Lund cohorts, respectively). In the second step, we used the stochastic oscillator model (**Figure 1A**) to relate the empirical peak location to the approximate age of the tumor. Because this step requires knowledge about the unknown stochastic (de-)methylation rate, we constrained the overall calendar age distribution across the Lund and TCGA cohorts to have a median of 3 years, which corresponds to the mean sojourn time in breast cancer.^4,47^ See Supplementary Methods for details.

### Purity adjusted analyses

Acknowledging the correlation between the epigenetic clock index *c_β_* and tumor purity, we derived a purity-adjusted epigenetic clock index 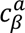 and repeated relevant correlation analyses with 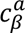 instead of *c_β_*. Because the epigenetic clock index was derived from the distribution of *β*-values of fCpG sites, we performed the purity adjustment at the level of *β*-values. For this, we assumed that the measured *β*-value at site 𝔦 (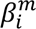) could be decomposed as a weighted sum of *β*-values of the tumor (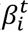) and the immune component (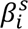),

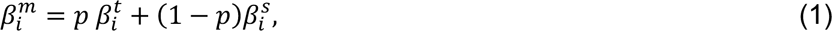

where *p* is the sample purity as measured by CPE. To estimate 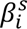 we combined the CIBERSORTx decomposition of the stroma (see section *CIBERSORTx*) with *β*-values of its constituent cells (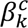) to obtain

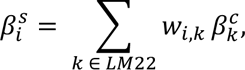

where *w_i,k_* is the fraction of cell type *k* (in the LM22 signature) in tumor sample *i*. The 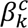 were estimated using published cell-type specific methylation values.^48^ Finally, the purity adjusted *β*-values were obtained by solving equation (1) for 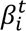 and truncating values below 0 and above 1 (necessary for <5.7% of the adjusted *β*-values).

### Statistical analyses

Correlations between two continuous variables were calculated using the Pearson correlation coefficient. The medians of continuous variables were compared using a two-sided Wilcoxon rank-sum test at significance level of 0.05. For each variable, tumors with missing values of that variable were excluded. All analyses and visualizations were performed in Python (3.9.19) and R (version 4.3).

## Code availability

All Python and R code used to produce the results in this paper are found on Github at https://github.com/danmonyak/EpiClockInvasiveBRCA (MIT License).

## ACKNOWLEDGMENTS

We gratefully recognize our funders who provided support for this work: National Institutes of Health (grant R01-CA271237 to M.D.R. and L.J.G.; grant U2C-CA233254 to E.S.H.; grant U54-CA217376 to D.S.), and Breast Cancer Research Foundation (grant BCRF-19-074 to E.S.H).

## COMPETING INTERESTS

The authors declare no competing interests.

## OVERVIEW OF SUPPLEMENTARY MATERIALS

**Supplementary Methods**: Mathematical modeling

**Supplementary Figure S1**: Purity-adjusted epigenetic clock index

**Supplementary Figure S2**: Pathologic correlates of the purity-adjusted epigenetic clock index

**Supplementary Figure S3**: Purity-adjusted pathway enrichment and immune decomposition analyses

**Supplementary Figure S4**: Sample correlations in the AURORA cohort

**Supplementary Table S1**: Associations of measured and adjusted clock indices with different markers

**Supplementary Table S2**: Associations of measured and adjusted clock indices with composition of the immune compartment (CIBERSORTx)

**Supplementary Table S3**: Subtype-specific distributions of clock index, mitotic age and calendar age

**Supplementary Table S4**: P-values of a two-sided Wilcoxon rank-sum test comparing the mitotic age distributions of each intrinsic subtype

## Notes

### Competing Interest Statement

The authors have declared no competing interest.

## REFERENCES

1 Duffy, S. W., Chen, H. H., Tabar, L. & Day, N. E. Estimation of mean sojourn time in breast cancer screening using a Markov chain model of both entry to and exit from the preclinical detectable phase. Sta$s$cs in medicine 14, 1531–1543 (1995).

2 Michaelson, J. et al. Estimates of breast cancer growth rate and sojourn time from screening database information. Journal of Womens Imaging 5, 11–19 (2003).

3 Shapiro, S., Goldberg, J. D. & Hutchison, G. B. LEAD TIME IN BREAST CANCER DETECTION AND IMPLICATIONS FOR PERIODICITY OF SCREENING1. American Journal of Epidemiology 100, 357–366 (1974). 10.1093/oxfordjournals.aje.a112046

4 Shen, Y. & Zelen, M. Screening sensitivity and sojourn time from breast cancer early detection clinical trials: mammograms and physical examinations. Journal of Clinical Oncology 19, 3490–3499 (2001).

5 Weedon-Fekjær, H., Vatten, L. J., Aalen, O. O., Lindqvist, B. & Tretli, S. Estimating mean sojourn time and screening test sensitivity in breast cancer mammography screening: new results. Journal of Medical Screening 12, 172–178 (2005). 10.1258/096914105775220732

6 Hannum, G. et al. Genome-wide methylation profiles reveal quantitative views of human aging rates. Molecular cell 49, 359–367 (2013).

7 Horvath, S. DNA methylation age of human tissues and cell types. Genome biology 14, 1–20 (2013).

8 Yang, Z. et al. Correlation of an epigenetic mitotic clock with cancer risk. Genome biology 17, 1–18 (2016).

9 Youn, A. & Wang, S. The MiAge Calculator: a DNA methylation-based mitotic age calculator of human tissue types. Epigenetics 13, 192–206 (2018). 10.1080/15592294.2017.1389361

10 Zhu, T., Tong, H., Du, Z., Beck, S. & Teschendorff, A. E. An improved epigenetic counter to track mitotic age in normal and precancerous tissues. Nature Communications 15, 4211 (2024). 10.1038/s41467-024-48649-8

11 Zhou, W. et al. DNA methylation loss in late-replicating domains is linked to mitotic cell division. Nature Genetics 50, 591–602 (2018). 10.1038/s41588-018-0073-4

12 Teschendorff, A. E. A comparison of epigenetic mitotic-like clocks for cancer risk prediction. Genome Medicine 12, 1–17 (2020).

13 Gabbutt, C., et al. Fluctuating methylation clocks for cell lineage tracing at high temporal resolution in human tissues. Nature biotechnology 40, 720–730 (2022).

14 Gabbutt, C., et al. Evolutionary dynamics of 1,976 lymphoid malignancies predict clinical outcome. medRxiv, 2023.2011.2010.23298336 (2023).

15 Teschendorff, A. E. On epigenetic stochasticity, entropy and cancer risk. Philosophical Transactions of the Royal Society B 379, 20230054 (2024).

16 Koboldt, D. C. et al. Comprehensive molecular portraits of human breast tumours. Nature 490, 61–70 (2012). 10.1038/nature11412

17 Johnson, K. C., Houseman, E. A., King, J. E. & Christensen, B. C. Normal breast tissue DNA methylation differences at regulatory elements are associated with the cancer risk factor age. Breast cancer research 19, 1–11 (2017).

18 Lewis, C. M. et al. Promoter hypermethylation in benign breast epithelium in relation to predicted breast cancer risk. Clinical Cancer Research 11, 166–172 (2005).

19 Shames, D. S. et al. A genome-wide screen for promoter methylation in lung cancer identifies novel methylation markers for multiple malignancies. PLoS medicine 3, e486 (2006).

20 Holm, K. et al. An integrated genomics analysis of epigenetic subtypes in human breast tumors links DNA methylation pàerns to chromatin states in normal mammary cells. Breast Cancer Research 18, 1–20 (2016).

21 Luo, Y. et al. Regional methylome profiling reveals dynamic epigenetic heterogeneity and convergent hypomethylation of stem cell quiescence-associated genes in breast cancer following neoadjuvant chemotherapy. Cell & Bioscience 9, 16 (2019). 10.1186/s13578-019-0278-y

22 Danielsen, H. E., Pradhan, M. & Novelli, M. Revisiting tumour aneuploidy — the place of ploidy assessment in the molecular era. Nature Reviews Clinical Oncology 13, 291–304 (2016). 10.1038/nrclinonc.2015.208

23 Ricke, R. M., van Ree, J. H. & van Deursen, J. M. Whole chromosome instability and cancer: a complex relationship. Trends in genetics 24, 457–466 (2008).

24 Chia, S. K. et al. A 50-gene intrinsic subtype classifier for prognosis and prediction of benefit from adjuvant tamoxifen. Clinical cancer research 18, 4465–4472 (2012).

25 Wallden, B. et al. Development and verification of the PAM50-based Prosigna breast cancer gene signature assay. BMC medical genomics 8, 1–14 (2015).

26 Rakha, E. A. et al. Breast cancer prognostic classification in the molecular era: the role of histological grade. Breast cancer research 12, 1–12 (2010).

27 Carter, C. L., Allen, C. & Henson, D. E. Relation of tumor size, lymph node status, and survival in 24,740 breast cancer cases. Cancer 63, 181–187 (1989).

28 Lolus, L. V., Amend, S. R. & Pienta, K. J. Interplay between Cell Death and Cell Proliferation Reveals New Strategies for Cancer Therapy. International Journal of Molecular Sciences 23, 4723 (2022).

29 Hanahan, D. & Weinberg, R. A. Hallmarks of cancer: the next generation. cell 144, 646–674 (2011).

30 Newman, A. M. et al. Determining cell type abundance and expression from bulk tissues with digital cytometry. Nature biotechnology 37, 773–782 (2019).

31 Desmedt, C. et al. Abstract S6-2: Characterization of different foci of multifocal breast cancer using genomic, transcriptomic and epigenomic data. Cancer Research 72, S6–2-S6-2 (2012).

32 Reyngold, M. et al. Remodeling of the methylation landscape in breast cancer metastasis. PloS one 9, e103896 (2014).

33 Garcia-Recio, S. et al. Multiomics in primary and metastatic breast tumors from the AURORA US network finds microenvironment and epigenetic drivers of metastasis. Nature Cancer 4, 128–147 (2023). 10.1038/s43018-022-00491-x

34 Aran, D., Sirota, M. & Butte, A. J. Systematic pan-cancer analysis of tumour purity. Nature communications 6, 8971 (2015).

35 Welch, H. G. & Black, W. C. Overdiagnosis in cancer. Journal of the National Cancer Institute 102, 605–613 (2010).

36 Li, J. et al. Molecular differences between screen-detected and interval breast cancers are largely explained by PAM50 subtypes. Clinical Cancer Research 23, 2584–2592 (2017).

37 Maley, C. C. et al. Classifying the evolutionary and ecological features of neoplasms. Nature Reviews Cancer 17, 605–619 (2017).

38 Brigham, Hospital, W. s., 13, H. M. S. C. L. P. P. J. K. R., 25, G. d. a. B. C. o. M. C. C. J. D. L. A. & Ilya, I. f. S. B. R. S. K. R. B. B. B. B. R. E. T. L. J. T. V. Z. W. S. Comprehensive molecular portraits of human breast tumours. Nature 490, 61–70 (2012).

39 Fredlund, E. et al. The gene expression landscape of breast cancer is shaped by tumor protein p53 status and epithelial-mesenchymal transition. Breast cancer research 14, 1–13 (2012).

40 Sherman, M. E. et al. The Susan G. Komen for the Cure Tissue Bank at the IU Simon Cancer Center: a unique resource for defining the “molecular histology” of the breast. Cancer Prevention Research 5, 528–535 (2012).

41 Seal, R. L. et al. Genenames. org: the HGNC resources in 2023. Nucleic Acids Research 51, D1003–D1009 (2023).

42 Parker, J. S. et al. Supervised risk predictor of breast cancer based on intrinsic subtypes. Journal of clinical oncology 27, 1160–1167 (2009).

43 Gendoo, D. M. et al. Genefu: an R/Bioconductor package for computation of gene expression-based signatures in breast cancer. Bioinformatics 32, 1097–1099 (2016).

44 Mootha, V. K. et al. PGC-1α-responsive genes involved in oxidative phosphorylation are coordinately downregulated in human diabetes. Nature genetics 34, 267–273 (2003).

45 Subramanian, A. et al. Gene set enrichment analysis: a knowledge-based approach for interpreting genome-wide expression profiles. Proceedings of the National Academy of Sciences 102, 15545–15550 (2005).

46 Bialic, M., Al Ahmad Nachar, B., Koźlak, M., Coulon, V. & Schwob, E. Measuring S-Phase Duration from Asynchronous Cells Using Dual EdU-BrdU Pulse-Chase Labeling Flow Cytometry. Genes 13, 408 (2022).

47 Bhà, R., et al. Estimation of age of onset and progression of breast cancer by absolute risk dependent on polygenic risk score and other risk factors. Cancer 130, 1590–1599 (2024).

48 Hannon, E., et al. Assessing the co-variability of DNA methylation across peripheral cells and tissues: Implications for the interpretation of findings in epigene9c epidemiology. PLoS genetics 17, e1009443 (2021).

