## Supplementary Figures and Tables for "Mapping the Temporal Landscape of Breast Cancer Using Epigenetic Entropy"

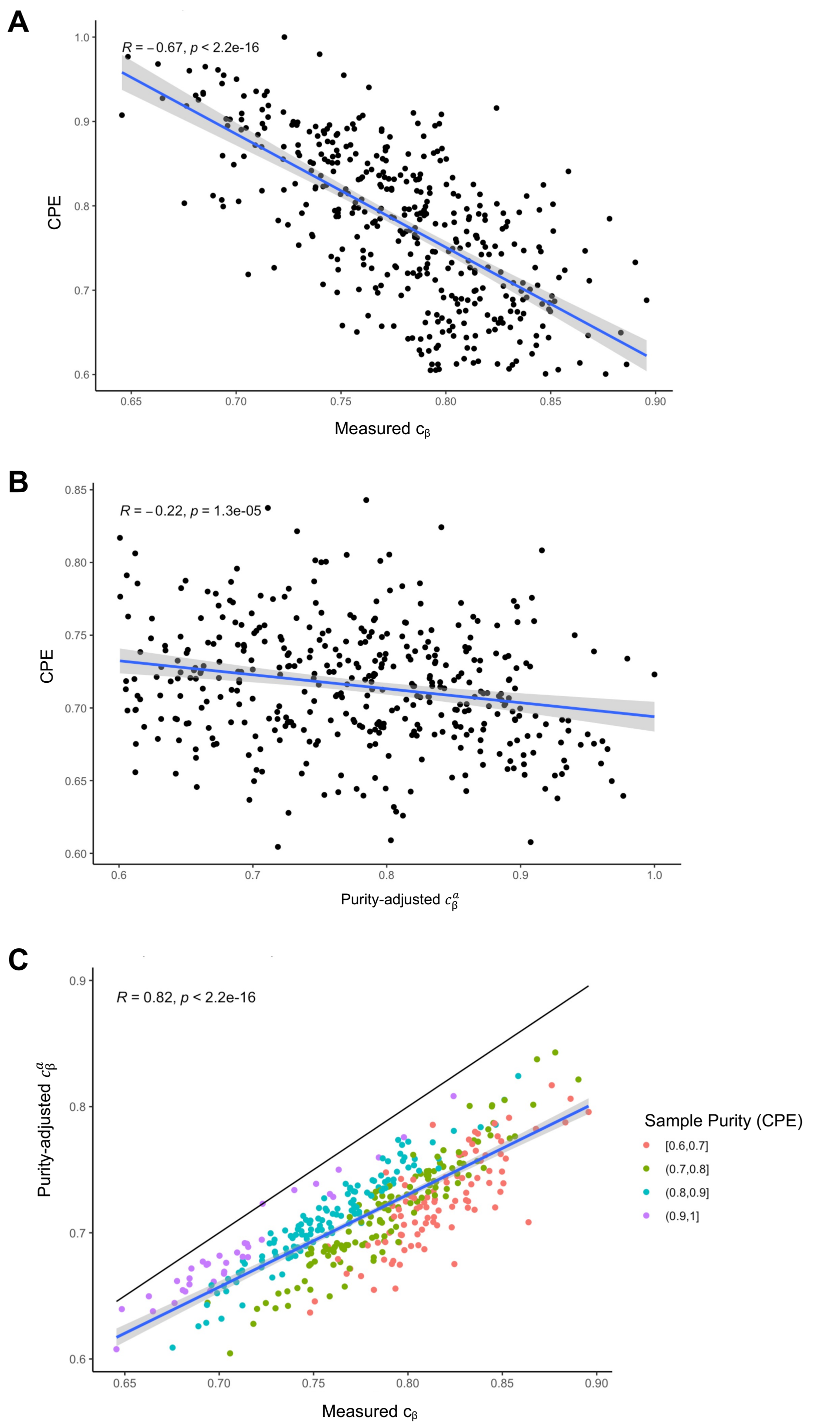


**Supplementary Figure S1. Purity-adjusted epigenetic clock index.** Data source: Invasive ductal carcinomas in TCGA, N=400. **(A)** Tumor purity, as measured by the consensus purity estimate (CPE), is negatively correlated with the epigenetic clock index $c_\beta$. **(B)** After adjusting the measured $\beta$-values for tumor purity, the resulting purity-adjusted epigenetic clock index $c_\beta^a$ was less correlated with CPE. **(C)** The purity-adjusted clock index was strongly correlated with, and invariably lower, than the measured clock index (solid line: identity line). R: Pearson correlation; blue line with gray band: linear regression line with 95% confidence band.

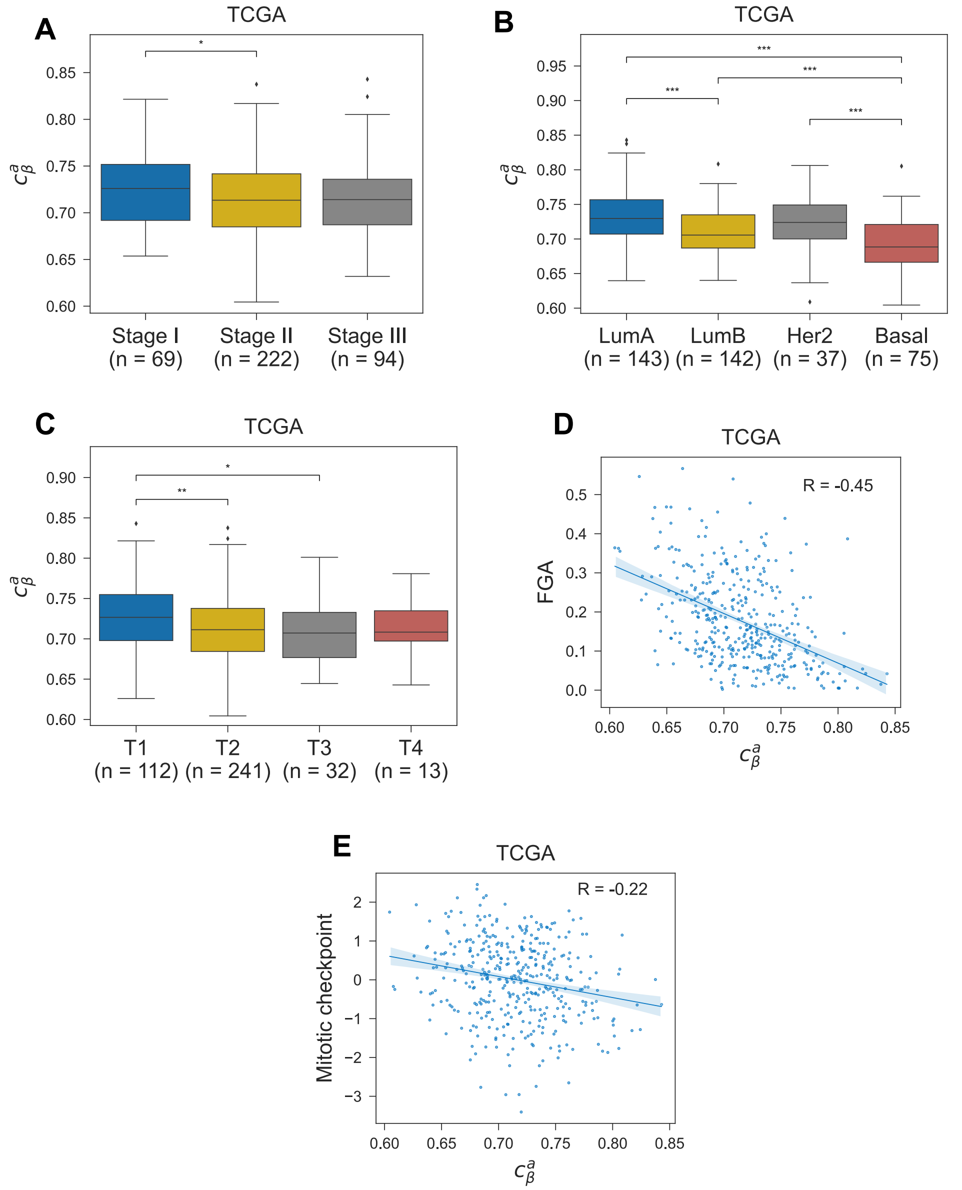


**Supplementary Figure S2. Pathologic correlates of the purity-adjusted epigenetic clock index.** The distribution of purity-adjusted epigenetic clock index values ($c_\beta^a$) among invasive ductal carcinomas in the TCGA cohort, by **(A)** tumor stage; **(B)** molecular subtype as predicted by the PAM50 algorithm; **(C)** T-stage; **(D)** fraction of genome altered (FGA) by copy number alterations; and **(E)** average expression of genes involved in M-phase and mitotic checkpoint regulation. Pairwise comparisons of medians were performed using a two-sided Wilcoxon rank-sum test (*P< 0.05, **P< 0.01, ***P< 0.001). Cohort size variations are due to the exclusion of tumors with missing values.

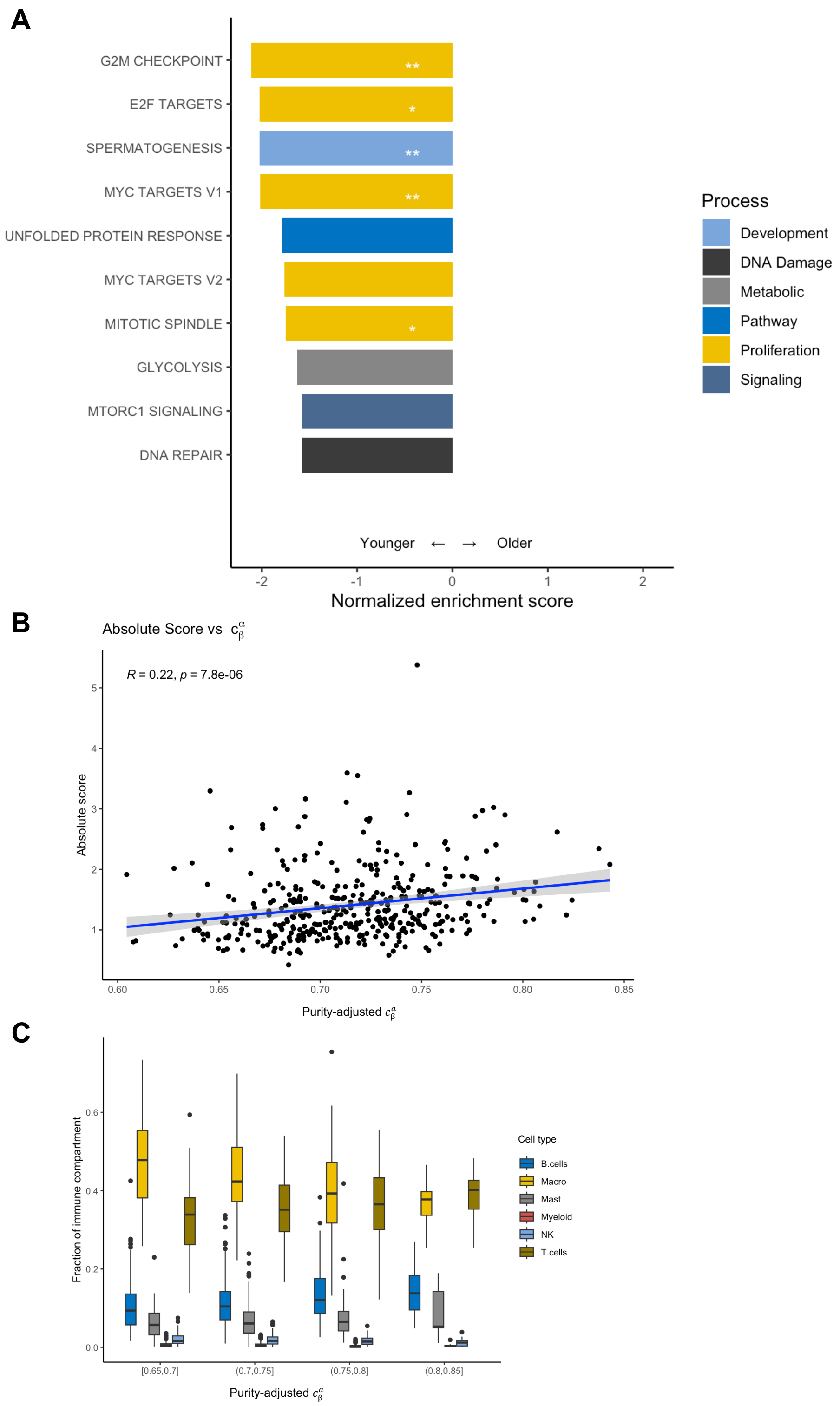


**Supplementary Figure S3. Purity-adjusted pathway enrichment and immune decomposition analyses. (A)** A gene set enrichment analysis (GSEA) was performed. Pathways with a negative enrichment score are enriched in younger tumors as measured by the purity-adjusted epigenetic clock index ($c_\beta^\alpha$). Only pathways with a false discovery rate (FDR) below 0.1 were included; *FDR<.05; **DFR<.01. **(B)** Purity-adjusted epigenetic clock index vs. the extent of immune infiltration (absolute immune score) as estimated by CIBERSORTx. **(C)** The immune compartment of each tumor was estimated using CIBERSORTx. The compartment fractions are shown for tumors of similar mitotic age, or epigenetic clock index ($c_\beta^\alpha$).

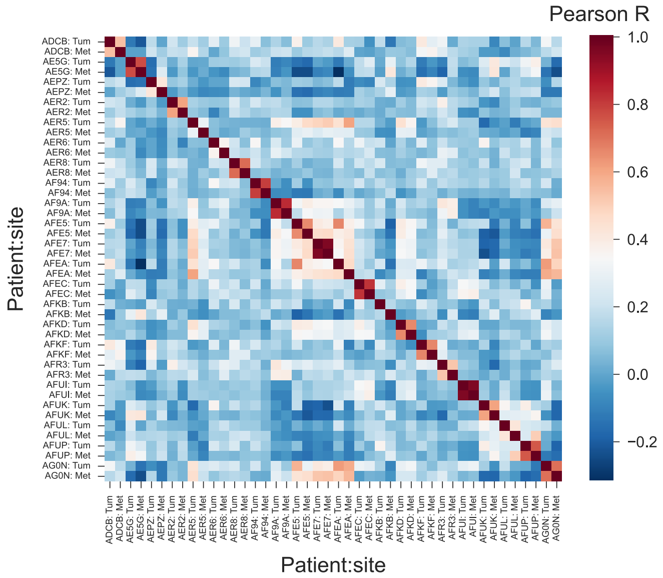


**Supplementary Figure S4. Sample correlations in the AURORA cohort.** In a cohort of 22 patients from the AURORA US Metastasis Project with paired primary tumor and metastasis samples, the $\beta$-values of the 500 fCpG sites are correlated within and between patients.

**Supplementary Table S1.** Associations of measured ($c_\beta$) and purity-adjusted ($c_\beta^\alpha$) clock indices with clinical, proliferation, and immune markers. FGA: fraction genome altered

| Correlate | Cohort | Measured clock index | | Adjusted clock index | |
|---|---|---|---|---|---|
| | | Pearson's R | P-value | Pearson's R | P-value |
| **Clinical markers** | | | | | |
| Age at diagnosis | TCGA | -0.18 | $2x10^{-4}$ | -0.19 | $2x10^{-4}$ |
| | Lund | -0.06 | $5x10^{-1}$ | NA | NA |
| Tumor size | Lund | -0.29 | $5x10^{-3}$ | NA | NA |
| **Proliferation markers** | | | | | |
| MKi67 | TCGA | -0.19 | $9x10^{-5}$ | -0.20 | $7x10^{-5}$ |
| MCM2 | TCGA | -0.19 | $3x10^{-5}$ | -0.19 | $2x10^{-4}$ |
| Mitotic checkpoint | TCGA | -0.24 | $1x10^{-6}$ | -0.22 | $1x10^{-5}$ |
| | Lund | -0.43 | $8x10^{-6}$ | NA | NA |
| S-phase fraction | Lund | -0.43 | $2x10^{-5}$ | NA | NA |
| FGA | TCGA | -0.56 | $4x10^{-34}$ | -0.45 | $4x10^{-21}$ |
| | Lund | -0.50 | $2x10^{-7}$ | NA | NA |
| **Immune markers** | | | | | |
| CD3D | TCGA | 0.37 | $1x10^{-14}$ | 0.24 | $1x10^{-6}$ |
| CD3E | TCGA | 0.44 | $2x10^{-16}$ | 0.30 | $1x10^{-9}$ |
| CD3G | TCGA | 0.41 | $2x10^{-16}$ | 0.27 | $3x10^{-8}$ |
| CD4 | TCGA | 0.42 | $2x10^{-16}$ | 0.19 | $2x10^{-4}$ |
| CD68 | TCGA | 0.11 | 0.031 | 0.02 | 0.66 |
| CD8A | TCGA | 0.34 | $1x10^{-12}$ | 0.23 | $4x10^{-6}$ |
| CD8B | TCGA | 0.12 | 0.018 | 0.07 | 0.14 |
| FOXP3 | TCGA | 0.33 | $2x10^{-11}$ | 0.16 | $1x10^{-3}$ |
| MS4A1 | TCGA | 0.32 | $4x10^{-11}$ | 0.30 | $5x10^{-10}$ |
| NCAM1 | TCGA | 0.03 | 0.54 | -0.01 | 0.87 |
| PTPRC | TCGA | 0.40 | $2x10^{-16}$ | 0.23 | $3x10^{-6}$ |

**Supplementary Table S2.** Associations of measured ($c_\beta$) and purity-adjusted ($c_\beta^a$) clock indices with extent (absolute immune score) and relative composition of the immune compartment (CIBERSORTx).

| Correlate | Measured clock index | | Adjusted clock index | |
|---|---|---|---|---|
| | Pearson's R | P-value | Pearson's R | P-value |
| Absolute immune score | 0.40 | $4x10^{-16}$ | 0.22 | $8x10^{-6}$ |
| **Relative composition** | | | | |
| B cells | 0.18 | $2x10^{-4}$ | 0.23 | $3x10^{-6}$ |
| Macrophages | -0.33 | $2x10^{-11}$ | -0.33 | $7x10^{-12}$ |
| Mast cells | 0.12 | 0.02 | 0.18 | $4x10^{-4}$ |
| Myeloid cells | -0.16 | 0.002 | -0.13 | 0.008 |
| Natural killer cells | -0.12 | 0.02 | -0.09 | 0.08 |
| T cells | 0.25 | $3x10^{-7}$ | 0.18 | $2x10^{-4}$ |

**Supplementary Table S3. Subtype-specific distributions of epigenetic clock index, mitotic age and calendar age.** The 25, 50, and 75 percentiles are shown for each subtype and for all tumors for each given metric. Only tumors with the middle peak located between 0.4 and 0.6 were included (see Main Methods).

| | $c_\beta$ | Mitotic age (Generations) | Calendar age (Years) |
|---|---|---|---|
| LumA | 0.8 (0.77, 0.83) | 168.65 (120.73, 225.37) | 6.51 (4.44, 9.61) |
| LumB | 0.77 (0.74, 0.79) | 126.62 (92.52, 155.8) | 2.36 (1.76, 3.27) |
| Her2 | 0.8 (0.77, 0.82) | 151.79 (120.83, 181.67) | 2.42 (1.41, 3.34) |
| Basal | 0.76 (0.73, 0.8) | 123.84 (90.23, 146.88) | 0.99 (0.62, 1.49) |
| Total | 0.78 (0.75, 0.81) | 139.95 (104.14, 181.7) | 3.0 (1.55, 5.6) |

**Supplementary Table S4. P-values of a two-sided Wilcoxon rank-sum test comparing the mitotic age distributions of each intrinsic subtype.** Only tumors with the middle peak between 0.4 and 0.6 were included (see methods).

| Mitotic Age | LumB | Her2 | Basal |
|---|---|---|---|
| LumA | $4x10^{-10}$ | $4x10^{-1}$ | $6x10^{-10}$ |
| LumB | | $4x10^{-5}$ | $4x10^{-1}$ |
| Her2 | | | $2x10^{-5}$ |
