## Supplementary Methods for "Mapping the Temporal Landscape of Breast Cancer Using Epigenetic Entropy"

#### *Measuring the Age of Individual Breast Cancers Using an Entropy-Based Molecular Clock*

Monyak et al., 2024

### 1 Estimation of tumor ages

To estimate tumor mitotic age based on the empirical distribution of $\beta$-values in the clock set of unbiased fluctuating CpG (fCpG) sites, we employ an analytically tractable, stochastic oscillator model of the (de-)methylation dynamics at the level of a single cell with two alleles.

We denote by $P(t) = (P_0(t), P_1(t), P_2(t))$ the time-dependent probability distribution of the fCpG site over the unmethylated ($P_0$), hemimethylated ($P_1$), and methylated ($P_2$) states, starting from the initial condition $P^0 = (p_0^0, p_1^0, p_2^0)$ at time $t = 0$. Since we are considering an unbiased fCpG site, the methylation and de-methylation rates are equal, or $\nu_{01} = \nu_{10} \equiv \nu$.

The forward Kolmogorov equation for this stochastic oscillator is

$$\begin{aligned}
\frac{dP_0}{dt} &= -2\nu P_0 + \nu P_1 \\
\frac{dP_1}{dt} &= 2(\nu P_0 - \nu P_1 + \nu P_2) \\
\frac{dP_2}{dt} &= \nu P_1 - 2\nu P_2.
\end{aligned} \tag{1}$$

After rescaling time by $\nu$, we get

$$\begin{aligned}
\frac{dP_0}{dt} &= P_1 - 2P_0 \\
\frac{dP_1}{dt} &= 2(P_0 - P_1 + P_2) \\
\frac{dP_2}{dt} &= P_1 - 2P_2,
\end{aligned} \tag{2}$$

which admits as analytic solution

$$\begin{aligned}
P_0(t) &= c_1 - c_2 e^{-2t} + c_3 e^{-4t} \\
P_1(t) &= 2c_1 - 2c_3 e^{-4t} \\
P_2(t) &= c_1 + c_2 e^{-2t} + c_3 e^{-4t}.
\end{aligned} \tag{3}$$

Next, we calculate the cell's expected methylation value for different initial conditions. For a cell starting in the unmethylated left "peak" state of the $\beta$-value distribution, that is with initial condition $P_A^0 = (1, 0, 0)$, the above solution becomes

$$\begin{aligned} P_0(t) &= \frac{1}{4} + \frac{1}{2}e^{-2t} + \frac{1}{4}e^{-4t} \\ P_1(t) &= \frac{1}{2} - \frac{1}{2}e^{-4t} \\ P_2(t) &= \frac{1}{4} - \frac{1}{2}e^{-2t} + \frac{1}{4}e^{-4t}. \end{aligned} \tag{4}$$

If we define $\beta_A(t)$ to be 0, 1, or 2 if at time $t$ the cell is hypo-, hemi-, or hyper-methylated, respectively, we find

$$\mathbb{E}\beta_A(t) = (0)P_0(t) + (\frac{1}{2})P_1(t) + (1)P_2(t) = \frac{1}{2}(1 - e^{-2t}). \tag{5}$$

Similarly, starting in the methylated state with initial condition $P_C^0 = (0, 0, 1)$, corresponding to the right peak of the $\beta$-value profile, the expected methylation value of the cell evolves as

$$\mathbb{E}\beta_C(t) = \frac{1}{2}(1 + e^{-2t}). \tag{6}$$

We can now draw from these results to estimate tumor age based on the empirical $\beta$-value distribution as follows. First, by decomposing the empirical distribution into left, central, and right peaks (see Main Methods for details on how to achieve this decomposition), we identify sets of fCpG sites that originated in the unmethylated, hemi-methylated, and fully methylated states in the founding tumor cell, respectively.

Next, denoting by $\hat{\beta}_A$ and $\hat{\beta}_C$ the respective models of the originally unmethylated and methylated peaks in the $\beta$-value distribution (see Main Methods), and assuming that the number of fCpG sites in each peak is sufficiently large, we can invoke the central limit theorem to approximate $\hat{\beta}_A(t) \approx \mathbb{E}\beta_A(t)$ and $\hat{\beta}_C(t) \approx \mathbb{E}\beta_C(t)$.

Based on equations (5) and (6), we can derive separate estimates of the (dimensionless) tumor age as

$$\hat{t}_A = -\frac{\ln(1 - 2\hat{\beta}_A)}{2} \tag{7}$$

and

$$\hat{t}_C = -\frac{\ln(2\hat{\beta}_C - 1)}{2}. \tag{8}$$

To recover dimensional estimates of tumor age, we note that the scaling factor is $\nu = \alpha\mu$ where $\alpha$ is the tumor's proliferation rate and $\mu$ is the per cell division and per allele probability of a (de-)methylation event. Therefore, tumor mitotic age $\tau$ corresponds to $t/\mu$ and tumor calendar age is obtained as $t/\alpha\mu$. In practice, we used the average of the two peaks' age estimates in (7) and (8).

Finally, while we were able to estimate tumor specific proliferation rates $\alpha_i$ (see Main Methods), the average (de-)methylation rate of fCpG sites is a priori unknown. For this reason, we set the median calendar age of tumors in the combined TCGA and Lund cohorts to 3 years (see Main Methods), and used this constraint to estimate the (de-)methylation probability ($\mu \approx 0.002$). Based on this estimate we then characterized the absolute distribution of tumor mitotic ages, as well as the relative distribution of tumor calendar ages (constrained to a median of 3 years).

#### 2 Estimation of proliferation rate from S-phase fraction

Our estimate of a tumor's proliferation rate $\alpha$ was based on the estimated S-phase fraction of cells $f$. To translate $f$ to $\alpha$ we used a simple model of cell cycle dynamics where the cell is either in state $S$ (in S-phase) or in state $\bar{S}$ (not in S-phase). Denoting by $r_1$ and $r_2$ the rates of transition from $S$ to $\bar{S}$ and vice-versa, the average times spent in $S$ and $\bar{S}$ are $T_S = 1/r_1$ and $T_{\bar{S}} = 1/r_2$, respectively.

In steady-state, the fraction of cells in S-phase is then given by

$$f = \frac{T_S}{T_{\bar{S}} + T_S} = \frac{\frac{1}{r_1}}{\frac{1}{r_1} + \frac{1}{r_2}} = \frac{r_2}{r_1 + r_2} \tag{9}$$

The proliferation rate $\alpha$ of a cell is the rate at which a cell completes the entire cell cycle, so it is the reciprocal of the time spent in both states once:

$$\alpha = \frac{1}{T_{\bar{S}} + T_S} = \frac{1}{\frac{1}{r_1} + \frac{1}{r_2}} = \frac{r_1 r_2}{r_1 + r_2} = f r_1 = f/T_S. \tag{10}$$
